# Metabarcoding, direct stomach observation and stable isotope analysis reveal a highly diverse diet for the invasive green crab in Atlantic Patagonia

**DOI:** 10.1101/2020.08.13.249896

**Authors:** Georgina Cordone, Mariana Lozada, Elisabet Vilacoba, Bettina Thalinger, Gregorio Bigatti, Darío A. Lijtmaer, Dirk Steinke, David E. Galván

## Abstract

The European green crab *Carcinus maenas* and its sister species *C. aestuarii* are highly invasive species causing damage to coastal ecosystems and contributing to severe economic losses worldwide. *C. maenas* was first detected at the Atlantic Patagonian coast in 2001. In this work, we studied the diet of the green crab in a recently invaded location in Golfo Nuevo, using three complementary techniques: direct stomach observation, metabarcoding of gut content and stable isotope analysis. Direct stomach observation and metabarcoding showed that green crabs have a broad omnivorous diet, ingesting most of the phyla present in the study area. Gut content metabarcoding allowed a detailed description of algal diversity and revealed other taxa that went unnoticed in the visual stomach analysis. Stable isotope analysis showed that the major contribution to the crabs’ diet was from the phytoplankton chain (by bivalve consumption) and not directly from algae. This study approach combining three complementary techniques also allowed us to detect some differences in the diet between sexes, which suggests that male and female crabs are not as ecologically equivalent as previously thought. Besides, we detected sequences corresponding to *C. aestuarii* suggesting that the green crab Patagonian population is a hybrid of both sister species. These findings are key to understanding the impacts green crabs can have on the local ecosystem.

## 1. Introduction

Invasive species, particularly invasive predators, are one of the main causes of worldwide species decline and biodiversity loss (Clavero and García-Berthou 2005; Doherty et al. 2016). Once arrived at a new location, they establish new ecological interactions with resident species and these new interactions can have positive or negative effects on native species or the entire native food web (David et al. 2017). The ways biological invasions impact food webs are complex and context-dependent as there are various direct and indirect mechanisms by which an invasive species can affect other taxa (Thomsen et al. 2011a, b). Preying upon a native species is a direct mechanism that usually ends up with the native species population decreasing in size, or even becoming extirpated (Bruno et al. 2005; David et al. 2017). Exploitation and apparent competition are indirect mechanisms that can also cause a reduction of native population sizes (White et al. 2006; David et al. 2017). On the other hand invaders can have positive effects for resident species. In fact, for species at higher trophic positions, invaders can increase food availability and create new habitats (Rodriguez 2006; Thomsen et al. 2010; Thomsen et al. 2014). The invaders’ impact depends on the food web’s invasibility which is ultimately related to network structure (Hui et al. 2016). Earlier work has shown that food webs with higher biodiversity, mainly understood as higher species richness, are more resistant to invasions; however, on the flip side, highly diverse communities generally support more invasive species (Fridley et al. 2007; Tilman et al. 2014; David et al. 2017). By using simulations, Romanuk et al. (2017) showed that trophic cascades are more frequent when invaders become secondary consumers with abundant prey and only few other predators. Consequently, if we want to accurately predict possible impacts of a newly arrived invasive species, we need to understand its trophic interactions within the invaded ecosystem.

Determining the diet of a particular species is a key requirement to understand its trophic ecology, but it is far from being a simple achievement. There are a series of conceptual and methodological aspects that must be taken into account (e.g. Traugott et al. 2013). It is not the same to document which organisms an animal ingests as listing which organisms are targets of a predator or a scavenger. Similarly, the ingestion of a certain item does not assure its digestion and assimilation. A variety of direct and indirect methods are currently used to analyze diet composition (Majdi et al. 2018; Nielsen et al. 2018). Methods include visual observation of feeding events and regurgitations, the visual inspection of faeces, pellets or stomach contents, and the molecular profiling of stomach contents or faeces using DNA sequencing or immunoassays. Each of these techniques has its advantages and disadvantages, which depend on the characteristics of the study, the geographic area under analysis, and the taxa that are being targeted. DNA-based techniques allow the characterization of feeding interactions at high taxonomic resolution and are very useful when visual recognition is not possible, and in cases of omnivorous species, where a mixture of DNA from different species is present in gut content. The most effective approach is metabarcoding (Taberlet et al. 2012; Pompanon et al. 2012; Clare 2014; Nielsen et al. 2018; Unger et al. 2020). Metabarcoding couples DNA barcoding (Hebert et al. 2003) with high-throughput sequencing of community samples (i.e. gut or fecal samples in the case of diet studies) and as a result it normally emerges as the superior technique for prey identification (Nielsen et al. 2018). In addition, it not only provides information on ingested organisms, but it also has the advantage of providing complementary information, e.g. on the gastrointestinal parasites of a predator. It should be borne in mind however, that metabarcoding success depends on various factors, including the availability of a relatively comprehensive database of target organisms (i.e. the prey items in the case of diet studies), the right choice of markers (to avoid low taxonomic resolution) and primers, and the possibility to deal with putative contamination as well as sequencing or bioinformatics errors (Braukmann et al. 2019). Moreover, it has two relevant limitations when used as the sole technique to study diet: it cannot detect cannibalism and it does not provide information on the developmental stage of the prey (i.e. whether eggs, larvae or adults have been ingested; Nielsen et al. 2018; Traugott et al. 2021).

Direct visual observation of stomach content is a traditional technique to identify diet composition, but its use is not recommended as the sole source of information, mainly because the digestion of prey material can severely limit detection and introduces biases towards certain taxonomic groups, such as shell-bearing organisms or arthropods that possess exoskeletal structures. Moreover, identifications largely depend on the skills of the taxonomists involved (Majdi et al. 2018; Traugott et al. 2021). However, direct observation can provide indispensable first-hand information, allows to quantify the ingested amount, and in certain cases it can distinguish between secondary predation and random ingestion (Hyslop 1980; Arnett and Whelan 2001; Zacharia et al. 2004). In addition, it can guide the DNA-based analysis and help solve some of the metabarcoding shortcomings mentioned above (such as the identification of the development stage of the prey or the presence of cannibalism). As a result, the combination of visual inspection and DNA-based techniques can be considered a wise choice to describe diet composition (e.g., Thalinger et al. 2018).

Aside from such direct methods, indirect tracers of diet composition are robust and commonly used tools. They include the characterization of biomarkers such as fatty acids, sterols or pigments present in consumer tissues and the determination of the isotopic compositions of C, N and S in specific tissues (e.g. Galloway et al. 2015; Stock et al. 2018). Stable isotope analysis (SIA) is based on predictable differences between the isotopic signature of an organism and its food resources. It allows for the quantitative estimation of fluxes in food webs, the determination of a species’ trophic level and insights into the patterns of resource allocation (Boecklen et al. 2011). In spite of these advantages, a potential confounding factor of SIA is the variability of trophic discrimination factors (TDF) and turnover rates, which may depend on environments, diet, taxon and tissue (Pinnegar and Polunin 1999; Philippsen and Benedito 2013; Hussey et al. 2014; Lefebvre and Dubois 2016). Similarly, isotope compositions are highly variable through space and time (Hyndes et al. 2013; Mackey et al. 2015). Therefore, these factors should be carefully considered when stable isotopes are used in diet studies.

These direct and indirect approaches are largely complementary as the former can identify the prey which is ingested, while the latter provide information on what is assimilated. Furthermore, indirect methods such as SIA provide information at longer time scales while DNA or visual analyses register consumption within short time periods (typically 6-48h) (Albaina et al. 2010; Hayden et al. 2014). In fact, recent trophic ecology studies successfully combined SIA and metabarcoding (Compson et al. 2019; Whitaker et al. 2019) and thereby started to fill a gap that has been previously highlighted for the field (Majdi et al. 2018).

The European green crab *Carcinus maenas* (Linnaeus, 1758) is an invasive species native to the North Atlantic Coast. To this day it has reached a worldwide distribution (Carlton and Cohen 2003). Not only *C. maenas* invaded new locations, but its sister species *C. aestuarii*, or hybrids of both, have also been detected at invaded sites (Darling 2011). These crabs are considered aggressive feeders due to their voracious predation on other invertebrates (McDonald et al. 2001). It has been shown that portunoid crabs are scavengers and predators, and that the green crab is an active carnivore feeding especially on mollusca and crustacea (Ebling et al. 1964; Elner 1981; Grosholz and Ruiz 1995; McDonald et al. 2001; Chen et al. 2004). Female and male green crabs are considered ecological equivalents as they do not show differences in their diets nor in their foraging behaviour (Spooner et al. 2007; Young and Elliot 2020). The green crab was first detected at the Patagonian coast in 2001, and twelve years later it was registered in Golfo Nuevo, 250 km north of the point of the first observation (Torres and González-Pisani 2016). The coast of Argentina currently bears a total of 129 introduced and 72 cryptogenic marine species, and it has been estimated that every 178 days a new invasion is detected (Schwindt et al. 2020). The interactions between the green crab and native as well as introduced species could have unpredictable consequences for the local trophic structure, as well as for ecosystem functioning. Very little is known about green crab ecology at the Argentinean coasts, in particular with regards to its local trophic interactions (Hidalgo et al. 2007; Young and Elliott 2020). Moreover, the observation that feeding behaviour is more aggressive in invasive than in native populations (Howard et al. 2018), leads to further uncertainties on how this predator could affect the invaded ecosystem.

In this study we aimed to shed light on the diet of the green crab in Atlantic Patagonian coastal environments by combining three techniques that have proven useful for diet tracing: visual stomach content inspection, stable isotope analysis, and amplicon sequencing of a gene fragment from stomach content DNA (metabarcoding). We addressed the following questions to understand possible impacts on the trophic structure of the affected coastal communities: What are green crabs eating in the coastal environments of Atlantic Patagonia? Are there differences in the diets of females and males in Atlantic Patagonia? Does the use of multiple techniques contribute to a more complete picture of the green crab’s diet?

## 2. Materials and Methods

### 2.1 Sample collection

Green crabs were collected at Punta Este beach in Golfo Nuevo, Patagonia, Argentina (42°46 ′ S, 64°57 ′ W). Sampling was done monthly from November 2018 to February 2019 (austral spring and summer) in shallow waters with depths of about 2-3 m at high tide. Crabs were captured manually by SCUBA diving and frozen at -20°C upon arrival to the laboratory both to kill them and correctly preserve them for subsequent analyses. A total of 223 specimens were captured. Individuals were separated into two groups: one for traditional visual analysis (n_(vis)_ = 183) and the other for metabarcoding (n_(met)_ = 40). Some of the individuals of each of these two groups (n_(sia)_ = 26 in total) were used for the SIA. The samples for the three techniques correspond to the same time period. Crab sizes were registered by measuring carapace width (CW) in cm. Subsequently, crabs were sexed and dissected. For specimens destined for visual analysis, stomach content was extracted and kept in 70% ethanol. For specimens destined for metabarcoding, stomach content was manually collected from frozen samples by letting them thaw and carefully extracting contents with a plastic pipette tip through the mouth of the crab. Contents were placed in 100% ethanol (Merck) and stored at -80°C until DNA extraction. Muscle from crab chela was extracted and frozen for SIA. In addition, five individuals of the mussel *Mytilus edulis* and five of the herbivorous snail *Tegula patagonica* were collected to establish isotopic baselines. This was done under the assumption that both the mussel and the herbivorous snail are primary consumers feeding mostly on micro- and macroalgae, respectively. We selected these two primary consumers as they integrate possible variations of environment isotopic values and their tissue usually shows shorter turnover rates than secondary consumers. As a consequence, primary consumers provide the most appropriate isotopic signatures for distinguishing between pelagic and benthic pathways in aquatic ecosystems (Post 2002).

### 2.2 Visual analysis of gut contents

The visual analysis of gut contents was done using a stereomicroscope (Zeiss Stemi, model 2000c). Prey was identified to the lowest possible taxonomic level following catalogues and keys of the local fauna (Fauchald 1977; Gosztonyi and Kuba 1996; Boschi et al. 1992; Spivak et al. 2019), as well as through expert consultation and by comparisons with fresh material. Presence of prey items in each stomach was recorded and the frequency of occurrence (FO_(vis)_%) calculated per prey item as the percentage of non-empty stomachs in which the item was recorded. For statistical analysis, we grouped prey into major groups (Annelida, Arthropoda, Mollusca, Chordata, Echinodermata and Algae). We used a bootstrap approach to estimate the FO confidence intervals for these groups. Bootstrap procedures are useful for non-parametric estimates and have been used in several studies of trophic ecology (Tirasin and Jorgensen 1999). This approach consists of re-sampling with replacement of the original data matrix and generates a new set of simulated matrices. The original matrix included green crab stomachs per row and prey items (presence or absences) in columns. Simulated matrices were generated by randomly choosing rows from the original matrix. This procedure was repeated until the simulated matrix reached the same number of rows as the original. Green crab stomachs could be chosen more than once per simulated matrix as resampling allowed for replacement. We simulated 1000 new matrices using the functions implemented in the R package ‘boot’ and calculated the FO for each prey item (Canty and Ripley 2021). This is based on the assumption that the frequency distribution of the simulated FO is unbiased and fully describes the distribution function of the parameter of interest (Efron and Tibshirani 1993; Mooney and Duval 1993).

### 2.3 Metabarcoding analysis of gut content

#### DNA extraction and COI amplification

All molecular analyses were carried out under clean-room conditions at the Centre for Biodiversity Genomics (University of Guelph, Canada). First, the ethanol of the gut content sample was evaporated in a fume hood overnight. Remaining pellets were lysed using 180µl ATL buffer + 20µl Proteinase K (20 mg/ml) per sample and incubated on a rocking platform at 56°C overnight. Afterwards, the DNA was extracted using the DNeasy Blood and Tissue Kit (Qiagen, Hilden, Germany), following manufacturers’ instructions except for an additional centrifugation step (14,000 rpm) for removing the buffer AW2 from the silica membrane and for elution in two steps using 100 µl of Buffer AE for each. A fume hood control and extraction control were processed along the stomach content samples to check for cross-contamination. DNA extracts were quantified using the Qubit dsDNA HS Assay Kit and a Qubit 3.0 Fluorometer (Thermo Fisher Scientific, MA, USA). For the amplification of gut content DNA, we used a two-step PCR protocol following Elbrecht et al. (2019). PCRs were carried out using the Multiplex PCR Master Mix Plus (Qiagen, Hilden, Germany), 0.5 mM of each primer (IDT, Skokie, Illinois), and molecular grade water (HyPure, GE, Utha, USA) for a total volume of 25 ml. For the first PCR step, amplification of cytochrome oxidase I gene was performed using primers BF3/BR2 (5’ CCHGAYATRGCHTTYCCHCG 3’ / 5’ TCDGGRTGNCCRAARAAYCA 3’) and 4 µl of DNA extract (BR2: Elbrecht and Leese 2017, BF3: Elbrecht et al. 2019; see Electronic supplementary material: Online Resource ESM 1). For the second PCR step, 1 µl of the product obtained from the first PCR was used for amplification using fusion primers with individual tags per sample (see Online Resource ESM 2, section 2.3) (Elbrecht and Leese 2017; Elbrecht et al. 2019; Elbrecht and Steinke 2019). Thermocycling conditions for both PCR steps included an initial denaturation at 95°C for 5 minutes, 24 cycles of denaturation at 95°C for 30 seconds, annealing at 50°C for 30 seconds, and extension at 72°C for 50 seconds; and a final extension at 68°C for 10 minutes. After the second PCR, amplification success and band strength were checked on 1.5% agarose gels. For several samples which showed no or only very weak amplification, both PCRs were repeated, but with 35 cycles instead of 24 in the first PCR. These replicates received individual tag combinations and were prepared for sequencing on an Illumina MiSeq platform along with the extraction controls and seven PCR negative controls.

Normalization of the second PCR products was performed using the SequalPrep Normalization Plate Kit (Invitrogen, CA, USA). Samples were subsequently pooled into one tube and cleaned using SPRIselect following the left side size protocol with a sample-to-volume ratio of 0.76X (Beckman Coulter,CA, USA). Library quantification was performed using the Qubit dsDNA HS Assay Kit and a Qubit 3.0 Fluorometer (Thermo Fisher Scientific, MA, USA). Sequencing was done at the University of Guelph (Advanced Analysis Centre) on an Illumina MiSeq using the 600 cycle Illumina MiSeq Reagent Kit v3 (2 × 300) and 5% PhiX spike in. Sequencing results were uploaded to the NCBI Sequence Read Archive (SRA, Genbank, accession: SRR13517441).

#### Sequence processing

Sequences were preprocessed using JAMP (https://github.com/VascoElbrecht/JAMP). Briefly, sequences were demultiplexed, and forward and reverse paired-end reads were merged. Primer sequences were removed, and sequences between 380bp and 440 bp (+/- 30bp of expected fragment length) were kept in the dataset, in order to preserve the most of the possible diversity present in the sample.

The resulting files were processed in two ways: i) the reads were quality filtered and taxonomy was assigned using the Multiplex Barcode Research And Visualization Environment in combination with the Barcode of Life platform (mBRAVE/BOLD; Ratnasingham 2019; Ratnasingham and Hebert 2007), and ii) quality filtering was carried out in JAMP and the resulting haplotypes were grouped into OTUs and blasted against the NCBI nucleotide database (for details see Online Resource ESM 2).

#### Description of crab diet based on molecular data

Taxonomic assignments obtained by the two pipelines (mBRAVE/BOLD-based dataset and JAMP/NCBI-based dataset) were manually curated. First, genus level taxonomic tables were screened for locally occurring species and, in the case of the mBRAVE dataset, the quality of the BOLD references (BINs) was determined to define the most reliable taxonomic level. For instance, genera which are not described for Argentina, not known from the marine environment, and/or whose BIN included multiple genera or could be considered unreliable (e.g. composed by only one sequence) were considered as “unidentified” members of that family/class. In addition, sequences identified as belonging to terrestrial arthropod taxa were removed prior to analysis due to contamination concerns. The NCBI dataset was curated at the OTU level, based on % similarity of the first hit to OTU representative sequences. Only hits with pairwise identity > 85% and alignment coverage > 75% were kept for analysis. Of these, representative sequences were assigned to: species (at least 98% similarity with the first hit), genus (95%), family (90%), and order (85%) (Elbrecht et al. 2017). For analysis of taxa occurrence, both analyses were combined into one that contained BOLD-based assignments plus NCBI assignments of taxa not detected in mBRAVE. The results were used to calculate the frequency of occurrence (FO_(met)_) at different taxonomic levels in the same way as FO_(vis)_ (see section 2.2). We decided to use occurrence data and not quantitative data such as read count because gut content material is usually at various states of digestion and the sequence counts are less likely to contain a meaningful quantitative signal compared to e.g. fecal matter (Nakahara et al. 2015; Deagle et al. 2019). In this way, we avoid incorporating possible biases due to the stochasticity of the amplification success and the variability in digestion times. Complete tables of taxonomic assignments are provided as Online Resources (ESM 3: Table S2 and ESM 4: Table S3).

### 2.4 Statistical analysis of diet description

To test if sample size was sufficient to obtain a representative description of the diet, we constructed cumulative trophic diversity curves using the quadrats methods following Koen Alonso et al. (2002) for both visual and metabarcoding datasets. We calculated trophic diversity as the Brillouin index (Hz) and considered diversity curves as asymptotic if the two previous values (i.e. n-1 and n-2) were in the range Hz ± 0.05 Hz for phylum and genus classification (Pielou 1966, Koen Alonso et al. 2002). Since this condition was met the number of stomachs in our study, n_(vis)_ and n_(met)_ was considered sufficient to describe the green crab diet.

To test differences between male and female green crab diet, we performed a distance-based multivariate Welch t-test (Wd* -test) both for Class-level assigned metabarcoding and visual analyses. The Wd* -test is a robust alternative to PERMANOVA when heteroscedasticity or imbalanced data are present (Alekseyenko et al. 2016; Hamidi et al. 2019). The tests were performed using a Bray-Curtis similarity matrix implemented in vegan and the MicEco R package (Oksanen et al. 2019; Russel 2020).

Alpha and beta diversity metrics, ordinations, and other statistical analyses were calculated using OTUs resulting from the JAMP routine. Analyses were performed using the R packages vegan and phyloseq (McMurdie and Holmes 2013; Oksanen et al. 2019). Briefly, OTU richness was calculated using the *rarefy* function in vegan, which gives the expected species richness in random community subsamples of a given size. A Kruskal-Wallis non-parametric test was used to analyze whether there were differences in prey richness between sexes. For the beta diversity analyses, a similarity matrix based on OTUs was built using Jaccard similarity index, from which NMDS ordinations were constructed.

### 2.5 Stable isotope analysis

Muscle samples for SIA were extracted from the chela of the frozen specimens and placed back in the freezer in individual tubes. We made sure that samples remained frozen during the entire procedure by using ice and sampling the crabs in small groups (4 individuals), as repeated freeze/thaw cycles can alter sample isotopic values (De Lecea et al. 2011). *M. edulis* and *T. patagonica* samples were obtained in the same way. After obtaining all the samples, they were dried for 48 h at 60 °C, ground into a fine powder and ∼1.25 μg per sample was sent to the Stable Isotopes Center at the University of New Mexico for C and N stable isotope determination. Internal standards used in the laboratory were blue grama, buckeye, cocoa powder, casein, tuna muscle, graphite, whey, elemental protein, serine and IAEA 1N. Standard deviations (SDs) of reference materials were 0.05‰ for δ^13^C and 0.07‰ δ^15^N. A total of 36 muscle samples were sent, of which 26 corresponded to green crabs (10 males, 11 females and 5 duplicates), five to *M. edulis* and five to *T. patagonica*. The error estimated through duplicate samples for green crabs muscle was 0.14‰ for both δ^13^C and δ^15^N values.

We estimated the trophic position (TP) of the green crabs using the two baselines (i.e δ^13^C and δ^15^N values of mussels and snails) and the Bayesian approach proposed by Quezada-Romegialli et al. (2018) (See equations in Online Resource ESM 2: eq. 1, eq. 2 and eq. 3). The TDF used for TP estimations were 1.80 ± 0.40 ‰ (δ^13^C) and 2.35 ± 1.26‰ (δ^15^N) following published values for omnivorous crustaceans (Parker et al. 1989; Vanderklist and Ponsard 2003; Rudnick and Resh 2005; Yokoyama et al. 2005). For TP estimations sex was taken into account (i.e. independent estimates were undertaken for female and male crabs). We performed pairwise comparisons of posterior TP as the probability of TP_A_ to be less than or equal to TP_B_ (TP_A_ <= TP_B_) for comparisons between sexes. We similarly performed pairwise comparisons of the proportional contribution of each baseline (α). The Bayesian probability was considered to be significant when p > 0.95 as an analogous of the frequentist p-value. Estimates of TP and α were obtained using the ‘TwoBaselinesFull’ model in the R package ‘tRophic Position’ (Quezada-Romegialli et. al. 2018).

The trophic niche of green crabs was described in terms of isotopic niche characteristics following the standard ellipse approach (Jackson et al. 2011). We calculated the standard ellipse area for males and females separately considering 40% of the data and corrected by sample size (SEAc) and the Bayesian standard ellipses area (SEAb) using the ‘SIBER’ and “rjags” packages on R (Jackson et al. 2011). Subsequently, we calculated the percentage of overlap between females and males SEA as the area at the interception divided by the SEA of male or female respectively. To test differences between SEAb we calculated 100,000 posterior iterations of SEAb based on our data set and estimated the probability that the bigger SEAb was higher than the smaller SEAb (i.e. we estimated the probability of male SEAb > female SEAb) (Jackson et al. 2011).

## 3. Results

### What are green crabs eating in the coastal environments of Atlantic Patagonia?

We considered the number of stomachs analyzed (n_(vis)_ =183 and n_(met)_ = 40) sufficient to provide a good description of the diet, because the trophic diversity curves were asymptotic and all (n-1) and (n-2) iterations were within (Hz – 0.05) range (Fig. S1 in Online Resource ESM 2). Overall, 15.32% of the visually analyzed stomachs were empty and 35.48% contained material that was impossible to identify due to its high degree of digestion.

Metabarcoding generated a total of 17.47 mio reads. After mBRAVE denoising and filtering 3,163,504 reads (259,167 ± 164,274 SD per sample) could be assigned to taxa, 2,390,785 (75,6%) of which matched *Carcinus*. After JAMP processing 1,984,045 quality filtered sequences (110,498 ± 80,004 SD per sample) were used for clustering and blasted against Genbank’s NR database. 1,125,343 of these were assigned to *Carcinus* (56,7%) (Table S4 in Online Resource ESM 6). After removing all matches to terrestrial arthropod taxa, a total of 727,116 reads were assigned to prey taxa. One sample did not provide any high-quality reads that could be matched to reference sequences with either the NCBI or BOLD database, and four samples did not contain any DNA that could be assigned to prey species (i.e. they only contained host/green crab DNA). The first sample was excluded from the analysis and the other four were considered as empty stomachs resulting in 10.2% of empty guts for metabarcoding.

The visual analysis of the gut content showed that green crabs at Golfo Nuevo waters had a broad diet in terms of phylogenetic diversity, with 23 different taxa identified (Table 1). Identifications were possible due to prey remains (typically hard parts) found in gut contents. Polychaetes were identified by jaws and mollusks by opercle and radula. We also found some well preserved organisms such as tanaidaceans and amphipods, but finding an almost fully intact organism was very unusual (see Fig. S2 in Online Resource ESM 1). Overall, we detected the presence of different prey taxa such as mollusks, algae, arthropoda, annelida, fishes and echinodermata. The main prey items were Mytilidae (bivalves) with a FO_(vis)_ of 35.6% followed by the gastropod *T. patagonica* (12.7%).

**Table 1.**
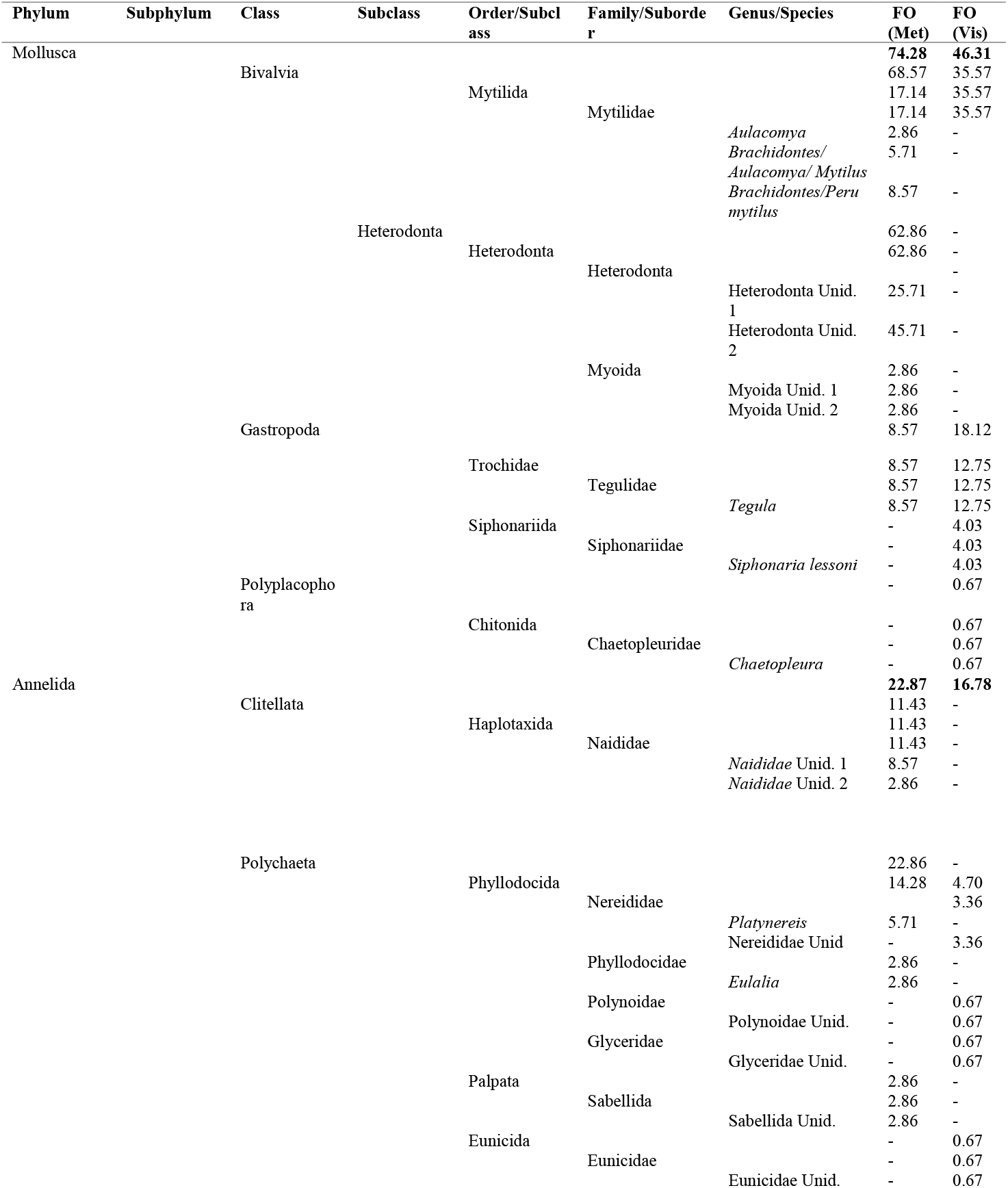
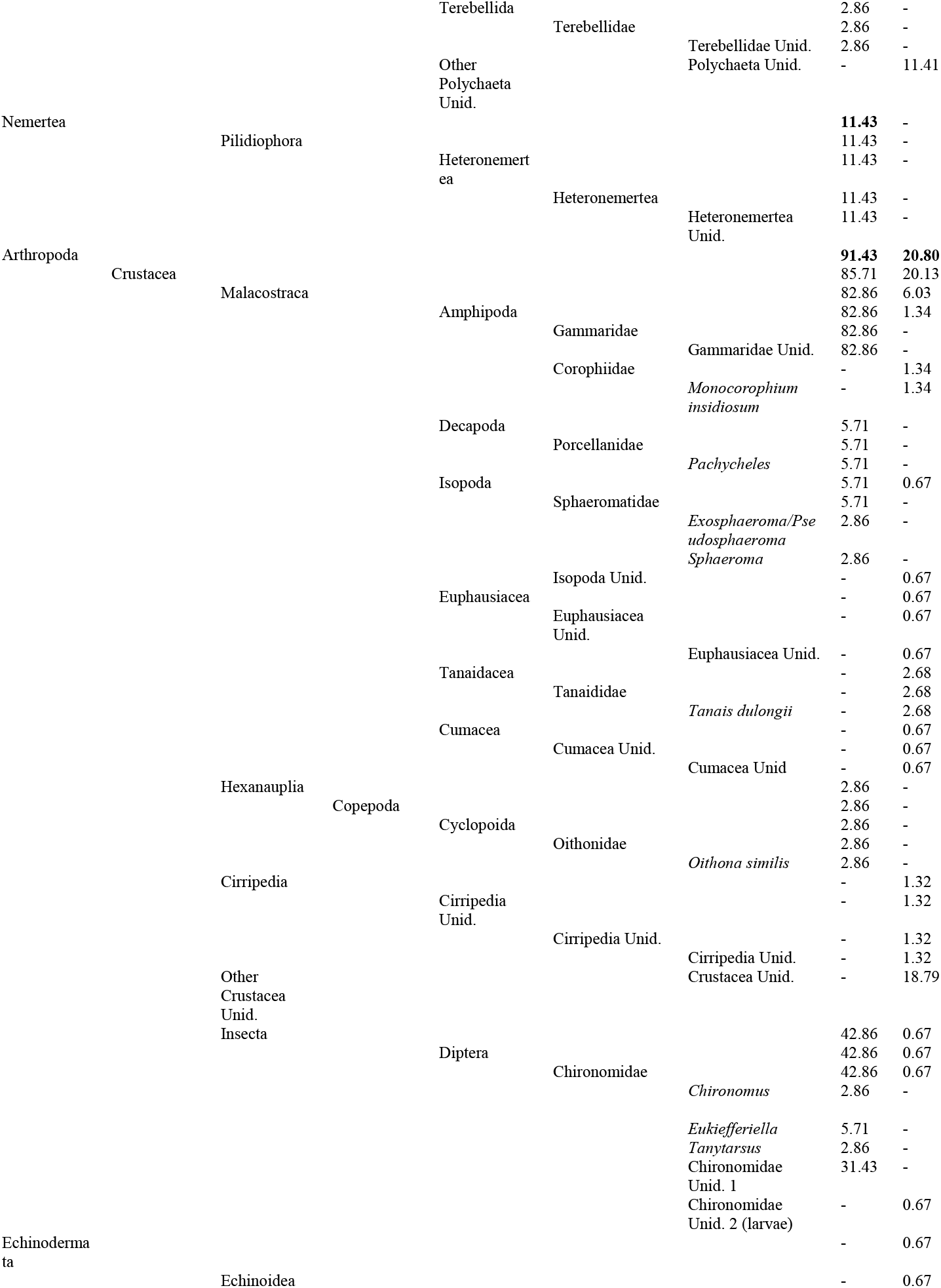
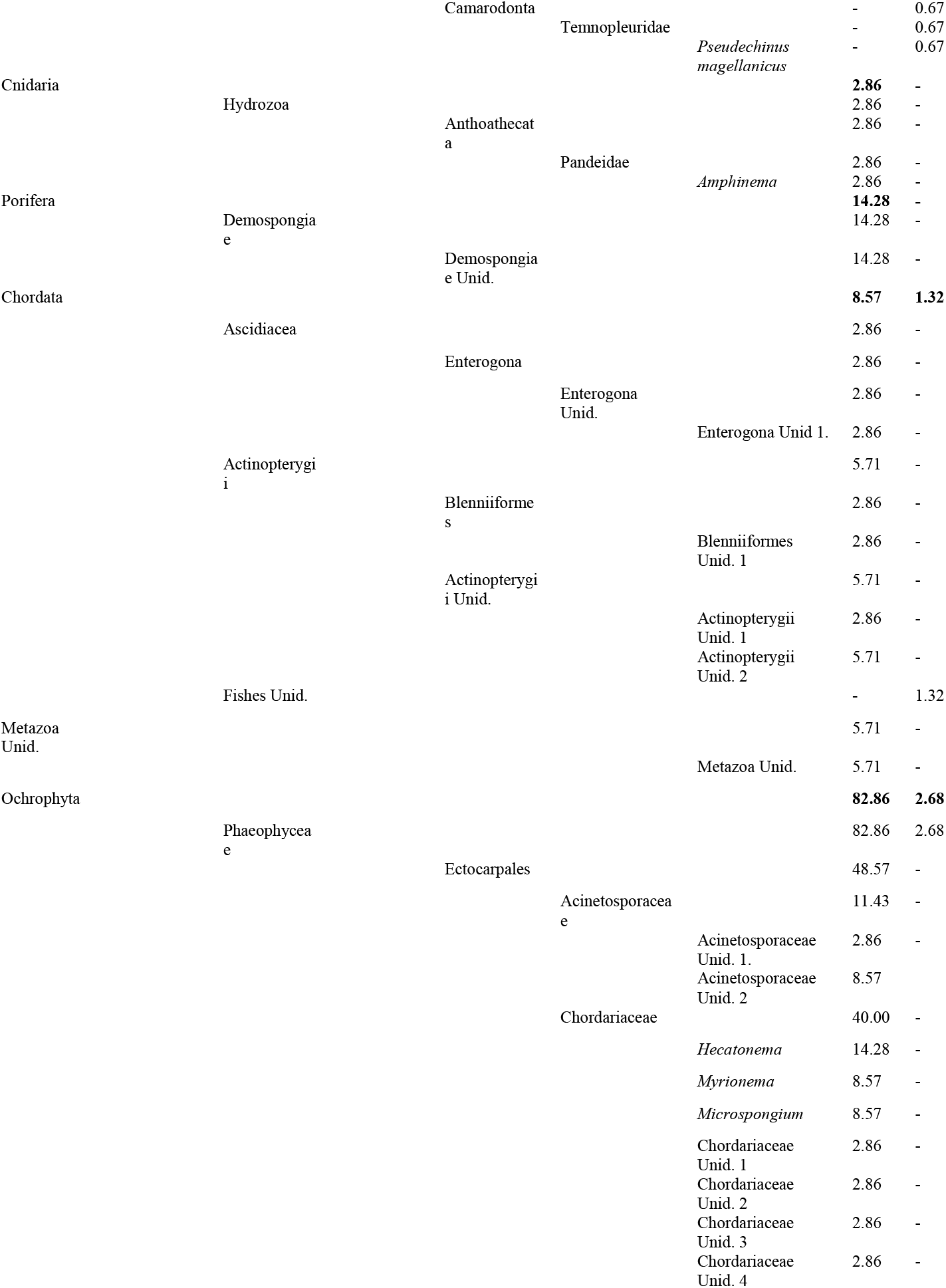
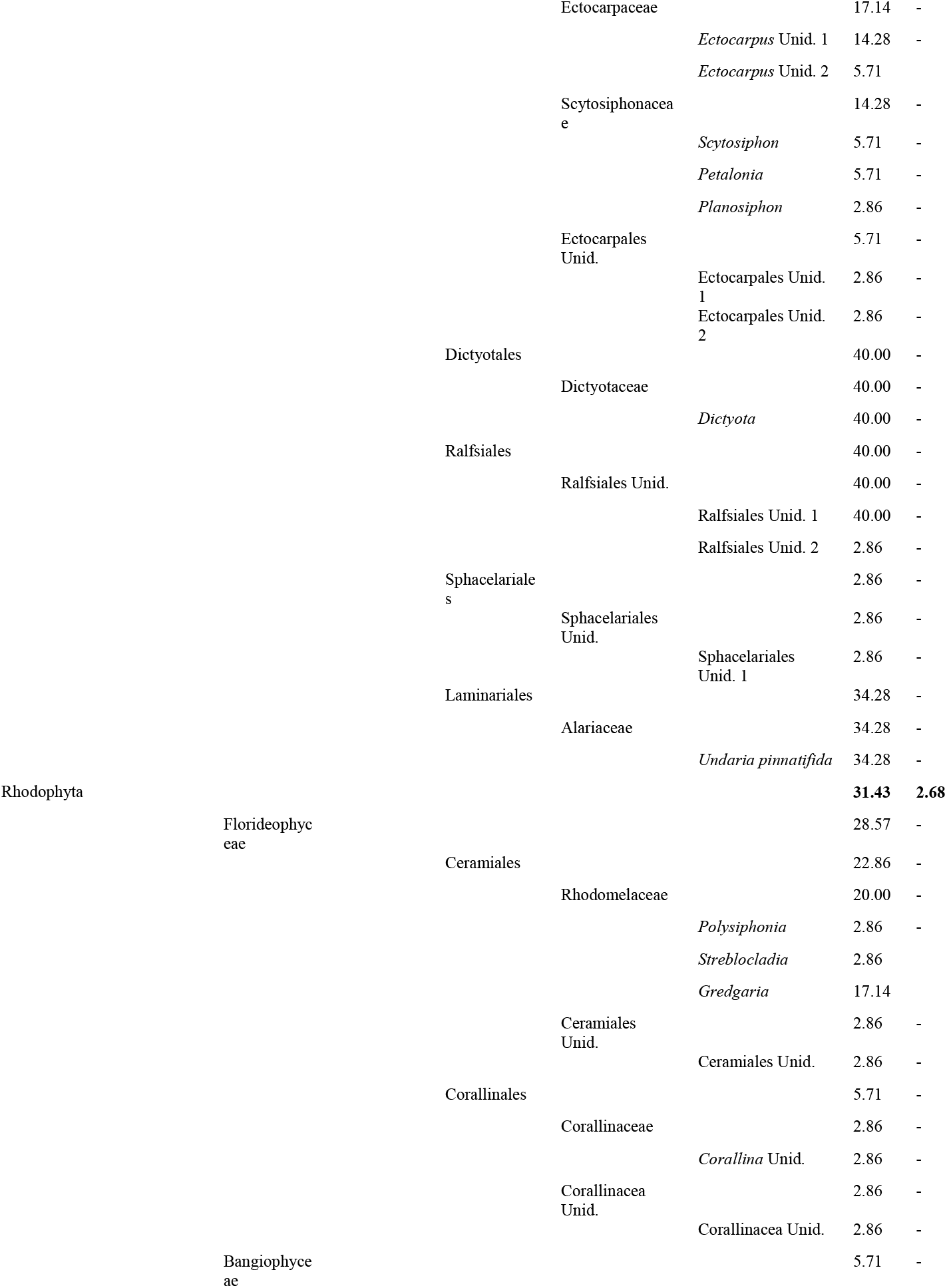
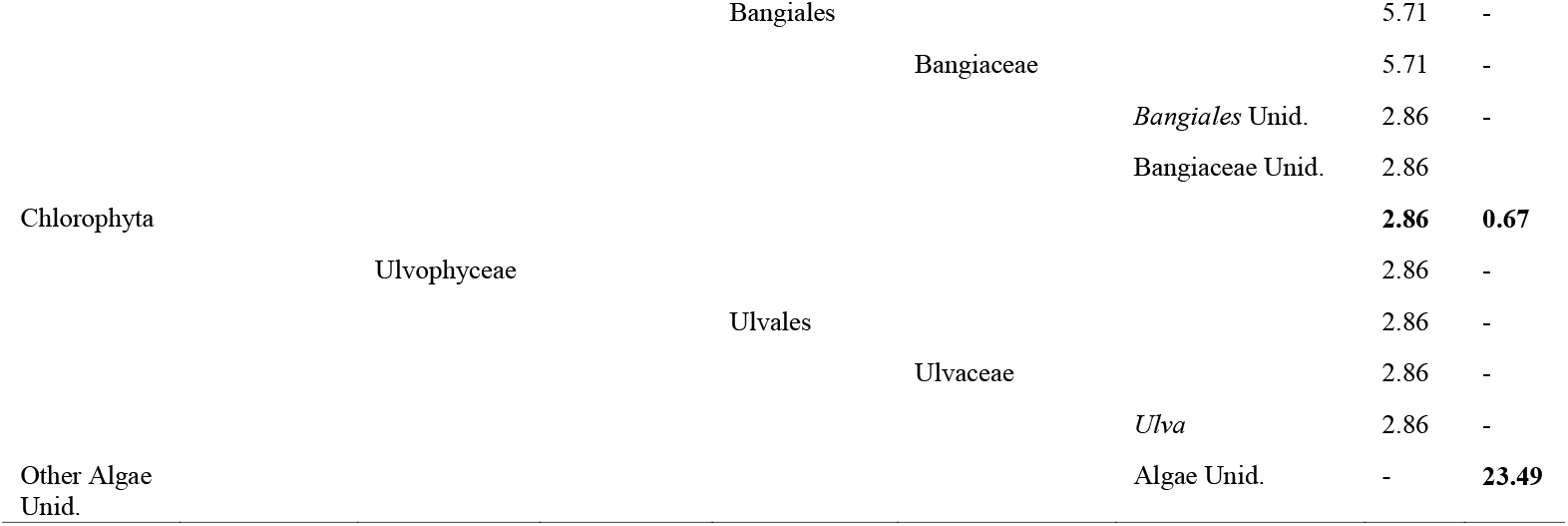
List of taxa in green crab stomach content determined by metabarcoding and visual analyses. Frequency of occurrence (FO%) for metabarcoding (met) and visual inspection (vis) of each taxa are shown.

Metabarcoding analysis also showed a very wide omnivorous diet, detecting a higher number of phyla and providing a better resolution at the genus level than the visual analysis. In general, FO_(met)_ were higher than FO_(vis)_, indicating that the level of detection with metabarcoding was higher than through traditional analysis. Here, Arthropoda was the dominant phylum in the crab’s gut, with a FO_(met)_ of 91.4%, followed by Ochrophyta (i.e. Phaeophyta) and Mollusca. The main prey items were amphipods of the Gammaridae family (82.9%), an unidentified species of Heterodonta (45.7%), and algae (*Dictyota* and an unidentified member of Ralfsiales, both 40%) (Table 1). The prey items with high FOs_(met)_ also had the highest read counts (see ESM 3 and ESM 4).

### Are there differences in diet between females and males in Atlantic Patagonia?

We observed small differences of prey items between males and females, evidenced by the overlapping of CIs of the simulated FO_(vis)_. The main difference was a higher frequency of arthropods and a lower frequency of molluscs in the female gut compared to male gut contents (Fig. 1a), but this difference was not significant (Wd*_(vis)_ = 3.14 and p-value_(vis)_ = 0.058). The FO_(met)_ showed fewer differences between male and female main prey groups (Fig. 1b); and we did not find significant differences between their diets either (Wd*_(met)_ = 0.85 and p-value_(met)_ = 0.66). Similarly, OTU ordinations showed a high degree of overlap between males and females (Fig. 2a).

**Figure 1.**
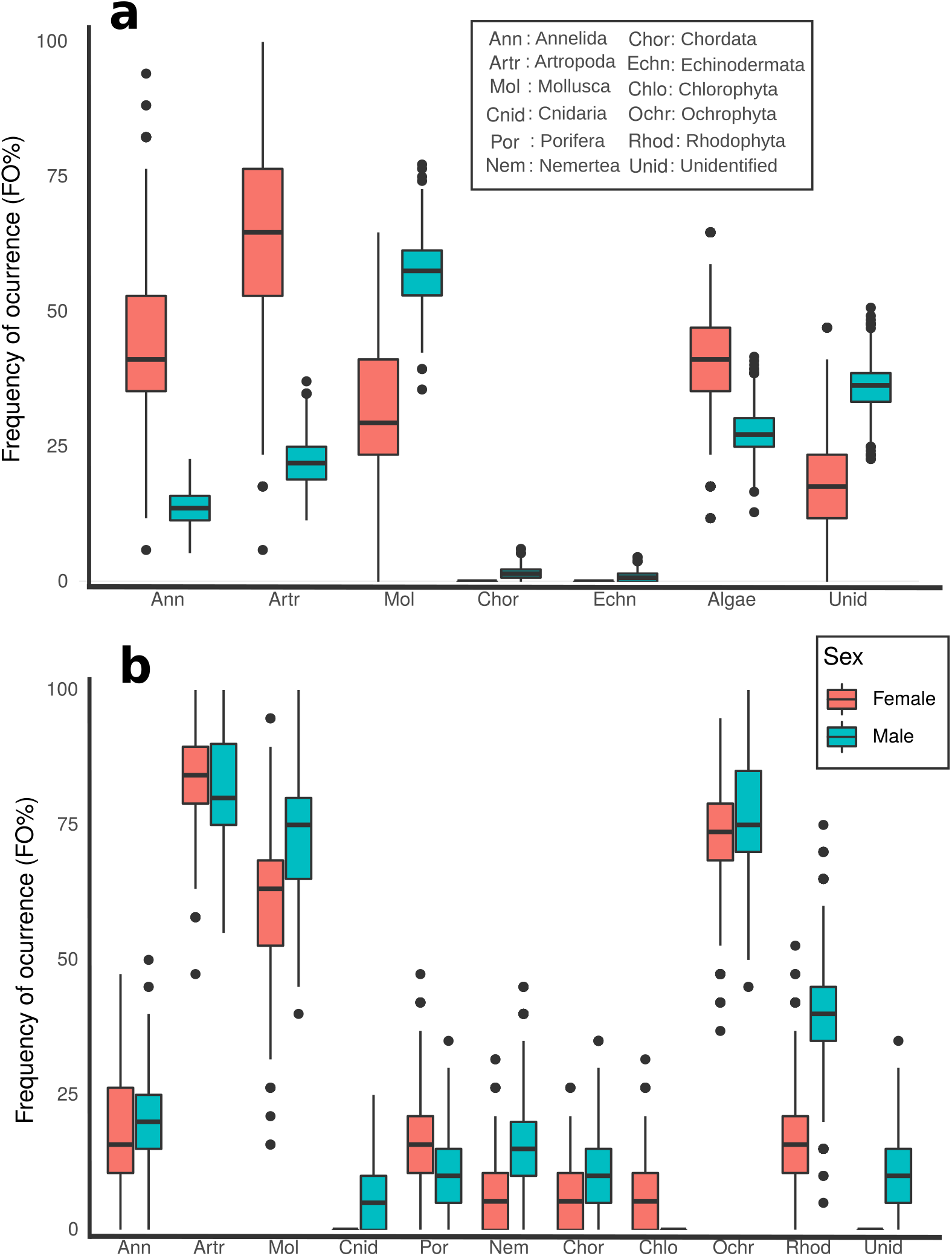
Frequency of occurrence (FO) boxplots of major prey groups in green crab diet from Patagonia, Argentina for: **a)** visual and **b)** metabarcoding analyses. The lower and upper hinges correspond to the first and third quartiles (the 25th and 75th percentiles). Points are outliers (data > 1.5* IQR) and horizontal lines are medians.

**Figure 2.**
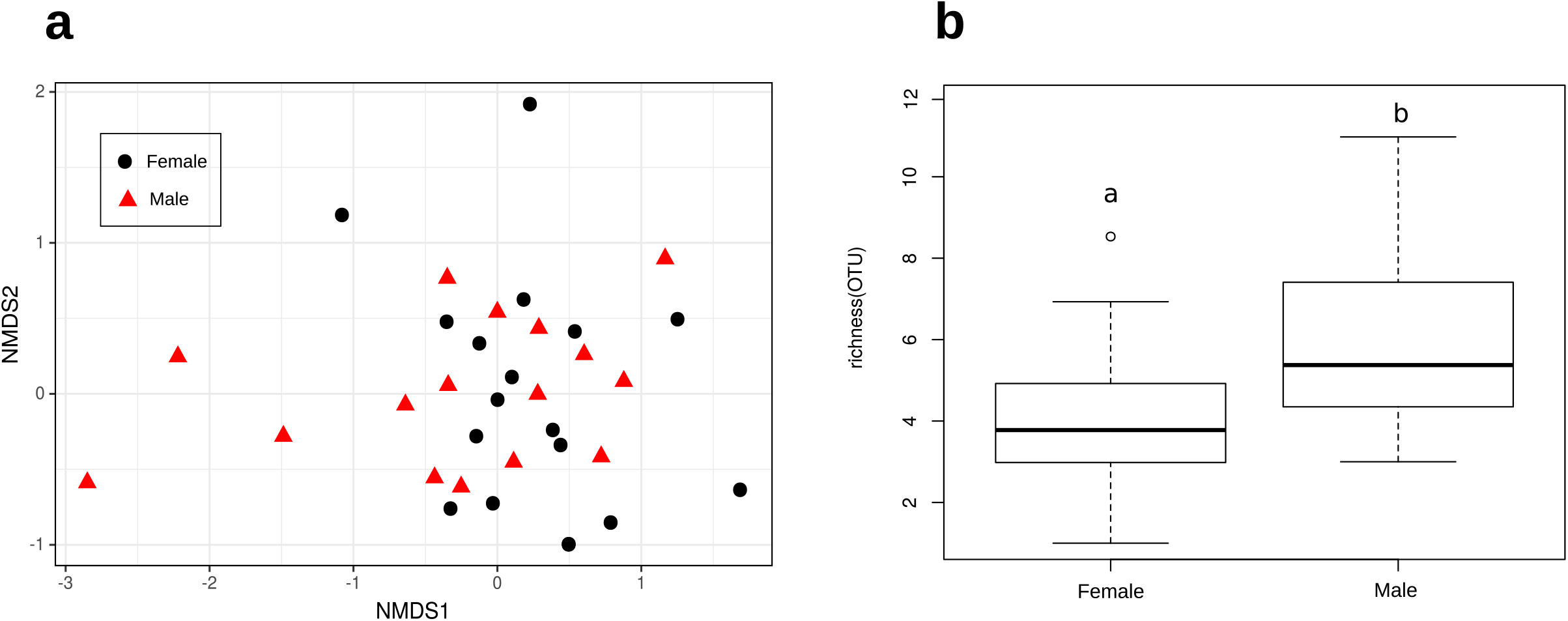
Differences between male and female diets in green crabs. **a)** Ordination of samples based on COI metabarcoding data (OTUs, NCBI-based assignments), depicted by sex. Distance measure: Jaccard index. **b)** Distribution of OTU richness (number of different OTUs) by sex. Letters “a” and “b” indicate statistically significant differences between sexes (p < 0.05; Kruskal-Wallis rank-sum test).

In spite of this small differentiation in prey items, we did detect differences between the gut contents of males and females in their trophic diversity. Metabarcoding OTU richness was significantly higher in males than in females (5.9 ± 2.3 and 3.9± 1.9, respectively; Kruskal-Wallis chi-squared = 6.6975, p-value = 0.009) (Fig. 2b). In fact, out of 64 OTUs considered prey, only 17 were shared between males and females (26%). These shared prey items corresponded roughly with the most frequently occurring OTUs, explaining the lack of clear grouping in ordinations (Fig. 2a). A great proportion of items (52%, 33 OTUs) were male-only OTUs while only 22% (14 OTUs) were female-only, indicating a higher richness in male diet.

Stable isotope analysis (SIA) results were in accordance with metabarcoding analysis. Males showed a broader trophic niche than females, with SEAc larger for males (SEAc_males_ = 1.03 and SEAc_females_ = 0.51, Fig. 4a). The Bayesian reconstruction of the SEA showed that this difference was significant (p = 0.05; Fig. 4b). However, it should be noted that some overlap at standard ellipses between males and females was present (11.89% of male SEA and 24.59% of female SEA) (Fig. 4a and Fig. S3 in Online Resource ESM 2). With respect to the trophic position (TP), female crabs showed a posterior estimated TP that was slightly higher than that for males (TP_males_ = 3.02, TP_females_ = 3.16 and TP_overall_ = 3.10, Fig 3a) but pairwise comparisons did not result in significant differences for TP or for alpha (α) posterior estimates (Table S5 in Online Resource ESM 2). The posterior estimates of the proportional contribution of each baseline showed the phytoplankton energy pathway to be more important than the macroalgae pathway for the nutrition of green crabs (α _males_ = 0.74, α _females_ = 0.88 and α _overall_ = 0.86, Fig 3b). This result is in accordance with bivalves (filter feeders) representing the main prey item.

**Figure 3.**
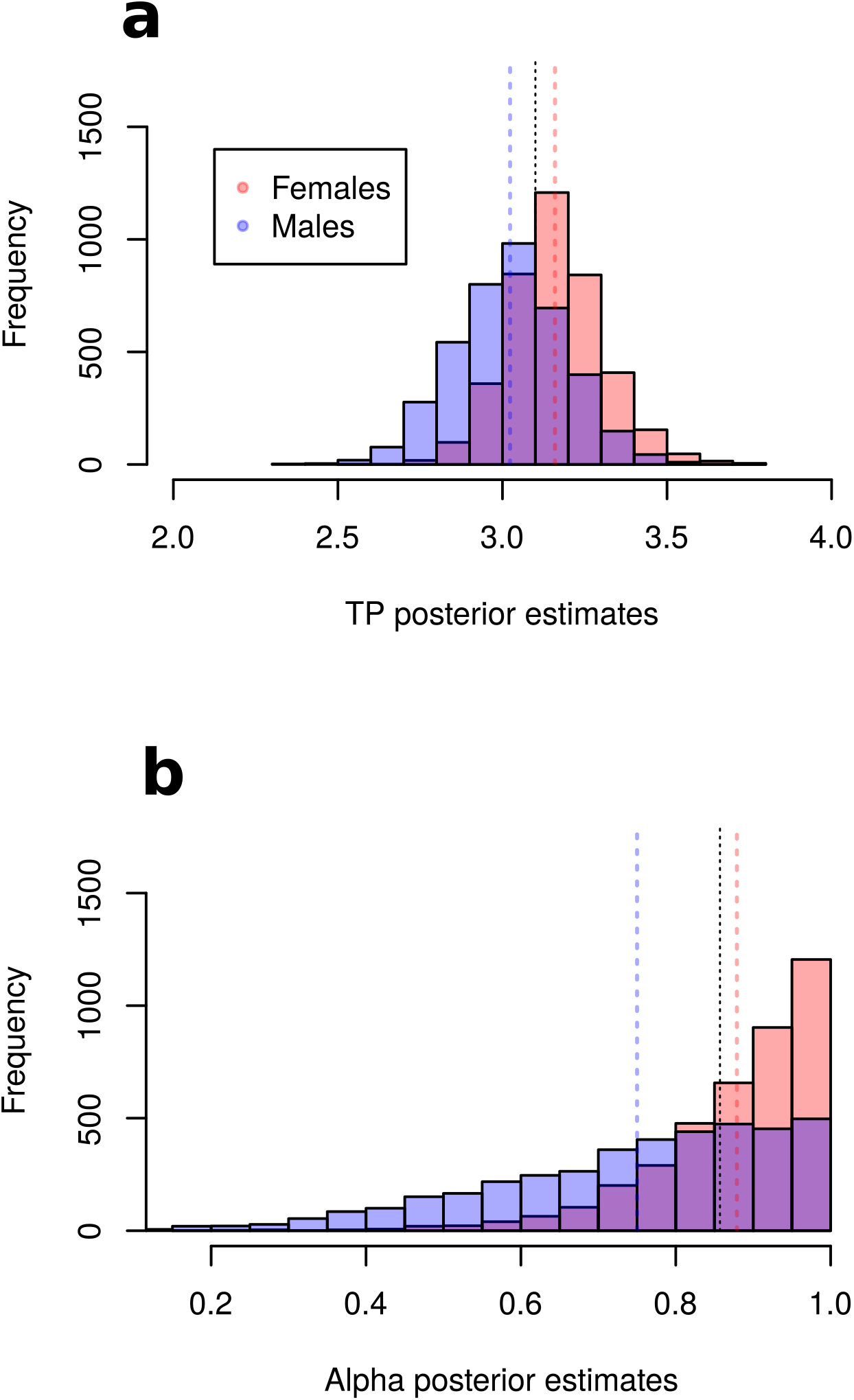
Posterior estimates from Stable Isotope Analysis of green crabs at Golfo Nuevo, Patagonia, Argentina. **a)** Trophic position (TP) for female and male green crabs **b)** Alpha (α) for female and male green crabs. Means are indicated by dotted lines (males: blue; females: pink; both sexes: black).

**Figure 4.**
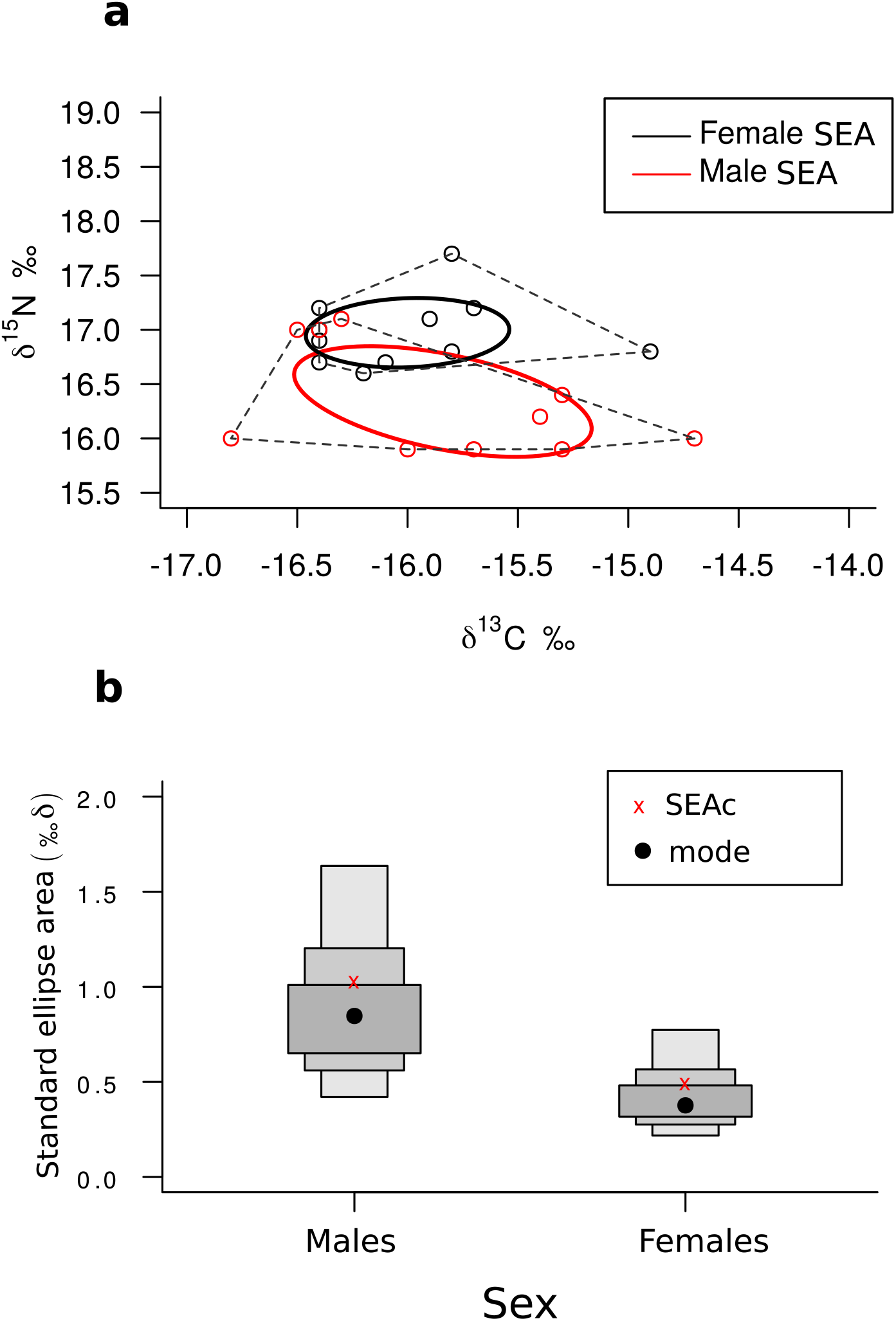
Stable Isotope Analysis of the green crabs at Golfo Nuevo, Patagonia, Argentina. **a)** δ^15^N‰ and δ^13^C‰ values for female and male green crabs. Ellipses represent the standard ellipses for each sex corrected for sample size. **b)** Standard ellipse area (‰2) estimated by Bayesian methods (SEAb) for male and female green crabs.

### Findings not linked to the green crab’s diet

Metabarcoding also showed other complementary results that are not diet-related but also relevant. In the first place, sequences from both *C. maenas* and *C. aestuarii* were detected in most samples, highlighting the possibility of the presence of a hybrid crab population in Patagonian coastal waters. Cannibalism could be an alternative explanation and, in fact, it has been previously observed in some green crab populations (Gehrels et al. 2017). However, cannibalism was ruled out because no parts of *C. maenas* were found in the visual inspections of the stomach contents. Besides, the high sequence abundance of the two species in each individual sample suggests hybridization (see Fig. S4 in Online Resource ESM 2). In addition, we detected sequences assigned to the genus *Carcinonemertes*, a nemertean parasite known to infect populations of this crab (Torchín et al. 1996).

## 4. Discussion

### What are green crabs feeding on in the coastal environments of Atlantic Patagonia?

Green crabs in the study site at the Patagonian Atlantic coast display a broad omnivorous diet, consuming the majority of phyla present at the study area. Phyla mostly reported for the sampling site (Mollusca, Arthropoda and Phaeophyceae, Rechimont et al. 2013) were also the prey items with the highest occurrences in our dietary samples. Rhodophyta and Annelida were also found frequently but at a lower occurrence rate. These results are consistent with previous studies in other areas, where green crabs have been classified as opportunist omnivorous, feeding on a wide range of prey species (Elner 1981; Singh 1991; Pardal et al. 2006). Other green crab populations, both native and invasive, have shown a strong preference for mollusks (bivalves and gastropods) (Young and Elliot 2020). In this study, mollusks were among the most common prey items based on visual examinations, whilst metabarcoding indicated species of amphipods of the Gammaridae family, which were present in more than 80% of the samples, as the most common prey item. Metabarcoding also revealed the presence of other prey items such as Nemertea, Porifera and Ascidiacea, which were not detected in visual analysis of stomach content.

Metabarcoding also provided a detailed description of algal diversity in the green crab diet. Although algae have been previously detected, identification achieved by visual analysis has been usually very poor (Cohen et al. 1995; Pardal et al. 2006; Wilcox and Rochette 2015). This is due to the little amount of intact algae that is usually found and the difficulties associated with taxonomic recognition. DNA analysis clearly represents an improvement for the identification of taxa that are notoriously difficult to recognize in degraded samples such as gut contents (Smith et al. 2005; García-Robledo et al. 2013; Taberlet et al. 2018). Although algae were frequently detected in our crabs’ diet, stable isotope analysis revealed that the major contribution to green crab diet stems from the phytoplankton food chain (i.e. the pelagic pathway of matter and energy). Green crabs in Atlantic Patagonia feed in an opportunistic manner, with bivalves and amphipods being their preferred prey; with those they also ingest some algae, but this does not significantly contribute to their nutrition.

### Are there differences in diet between female and male green crabs?

In this study, we found some differences between male and female green crab diets suggesting that the two sexes are not as ecologically equivalent as previously thought (Spooner et al. 2007; Young and Elliot 2020). Male and female crabs were found to have similar diets and trophic positions (TP), but males had a broader diet than females (i.e. higher trophic diversity). This difference was evidenced by both the isotopic niche and metabarcoding OTU richness. One possible explanation for males having a higher trophic diversity is related to differences in behavior between the sexes. Female crabs are usually found in deeper waters and spend a greater part of their life in the subtidal area (Reid et al. 1997; Klassen and Locke 2007). It is thought that green crabs migrate towards the intertidal zone to feed; then, females spending more time in the subtidal zone would have less opportunities to encounter rare prey than males, leading to a lower trophic diversity (Dare and Edwards 1981). Another possible explanation for the sex differences in trophic diversity is the sexual dimorphism of the chela. Male crabs have bigger and stronger master chela than females and chela size has been shown to be a good predictor of prey handling time and thus can influence feeding rate and energy intake (Elner 1980; Howard et al. 2018). In this context, if males can manipulate their food faster and thus spend less time on each prey item and more time searching for food, they could not only eat more, but also increase their chances of encountering less common prey items.

Finally, it should be noted that dietary overlap between the sexes remains high and that the differences are rather subtle. New dietary studies focusing on seasonality would be very helpful in exploring these differences in Atlantic Patagonia.

### What changes can green crabs trigger in local food webs?

Green crabs from newly established populations have shown to be superior foragers compared to long-established invaders (Rossong et al. 2012). Therefore, we expect that the most dramatic changes in local food webs will occur in the early stages of the invasion until a new equilibrium state with new relative abundances has been reached. However, such a new food web state can bring serious consequences for native populations and even lead to extirpation of some species. The potential impact of green crabs as predators at the study site can be illustrated by comparing results to an earlier survey that was part of the Census of Marine Life, NAGISA project. In this study, a total of 35 species of benthic invertebrates and 29 macroalgae were recorded (Rechimont et al. 2013), while in the present study we found 31 benthic invertebrates and 23 macroalgae in the stomachs of green crabs. Hidalgo et al. (2007) observed that green crabs are bigger and feed with greater voracity than native predators on Patagonia’s rocky shores. They have provoked negative impacts worldwide such as damaging populations of *M. edulis*, American oysters (*Crassostrea virginica*), and rock crabs (*Cancer irroratus*), among others. They have even caused major economic losses for different fisheries (DeGraaf and Tyrrel 2004; Miron et al. 2005; Sigurdsson and Rochette 2013; Leignel et al. 2014; Young and Elliot 2020). In addition, bird mass mortality has been attributed to the transmission of parasites by green crabs (Camphuysen et al., 2002). Coastal birds in Patagonia are already incorporating green crabs into their diet, according to a recent kelp gull (*Larus dominicanus*) study that showed green crab as one of their prey items (Yorio et al. 2020). Carcinophagus fish (e.g. *Mustelus schmitti*) are likely to incorporate green crabs into their diet as well, although this has not been confirmed yet (Alessandra Pasti pers. comm. July 2020). The fact that the nemertean parasite *Carcinonemertes* was observed in one of our samples highlights the possible risk of dispersion of these parasites, which had not been reported before among green crabs or native crabs in Argentina (Bigatti and Signorelli 2018).

Community structure at intertidal Patagonian rocky shores is driven by harsh physical conditions, rather than consumer pressure (Bertness et al. 2006; Hidalgo et al. 2007). Invasive green crabs could have a negative impact on the community structure not only through the increase in predation pressure, but also by interfering with processes that maintain biodiversity such as facilitation (Silliman et al. 2011). That means green crabs could trigger both direct and indirect effects on Patagonian food webs. We predict that changes will begin with their preferred prey: mollusks, especially bivalves, as well as crustaceans, and then scale up to the entire food web through a cascade effect (White et al. 2006; Davies et al. 2017). We also predict that high predation on foundational species such as mitylids will constitute a loss of refuge for other intertidal species, which could result in a decline in biodiversity (Hidalgo et al. 2007). Changes at Golfo Nuevo coasts have been recently observed such as a dramatic decrease of *Perumytilus purpuratus* and *Brachidontes rodriguezii* populations (Mendez et al. 2019); however, the causes for the mass mortality are unknown and it is currently not possible to establish whether it relates to the green crab invasion or to environmental factors. In addition, other species of common native crabs are much more difficult to find at the intertidal since the arrival of green crabs, suggesting that they could have a negative interaction with other species of local crabs.

### Does the use of multiple techniques contribute to a more complete picture of the green crab’s diet?

The inclusion of complementary techniques to trophic ecology such as stable isotope analyses and metabarcoding can add new results that would otherwise go unnoticed when only relying on traditional techniques such as visual inspection of gut contents. These techniques complement each other and our study adds to previous work that demonstrates the usefulness of combining SIA and metabarcoding (e.g. Compson et al. 2019; Whitaker et al. 2019). Metabarcoding analysis contributes to the knowledge of diet richness and provides quantitative information about it. Stable isotope analysis allows for estimates of species trophic levels and trophic pathways (i.e. which sources contribute the most to species nutrition). The combination of these two techniques in our study provided information on different aspects of trophic ecology, such as interspecific prey-predator interactions and intrapopulation differences. In particular, we observed differences between male and female diets related to niche breadth and we achieved a more detailed taxonomic resolution for certain taxa due to the use of metabarcoding.

It should be mentioned at this point that visual inspections were not redundant in our analysis. In addition to the fact that they helped to rule out the possibility of cannibalism (suggesting that this is a hybrid population of *C. maenas* and *C. aestuarii*), some prey items found in the visual inspections were not detected by metabarcoding (for example: *Siphonaria lessoni* or *Pseudechinus magellanicus*). We think that the taxa detected only by visual inspection represent real prey items (not false positives generated by visual inspection) as they (Mollusca, Annelida, Arthropoda, Echinodermata, Fishes) have been previously observed as prey items of other green crab populations along the world and they are conspicuous in the intertidal. We have also observed this in laboratory tests, e.g. for S*iphonaria lessoni*. In fact, the discrepancy between metab.arcoding and visual inspection is showing some limitations of metabarcoding in our case study: First, the primers that are used for metabarcoding are not perfectly universal, which means that some taxa are less likely to be amplified (mainly fish and algae, but also some mollusks are affected). In addition, some of the taxa that were present in the gut content are not part of the reference library yet. Finally, distinct hard parts of some taxa can withstand digestion and are easy to identify morphologically, even after all the DNA has been broken down during digestion. Therefore, visual inspection is still recommended and can yield results that complement those of molecular methods such as metabarcoding, particularly in geographic areas or particular environments that have not been heavily barcoded yet, such as the Atlantic Patagonian coast. However, it is important to note that the visual and metabarcoding techniques have different sensitivity to prey detection, so a strict comparison between the two techniques is not possible. In fact, the visual and metabarcoding data collected in this study were used in a complementary, non-comparative way.

In conclusion, the complementary techniques used in this study allowed us to better understand the trophic biology of the green crab in a recently invaded habitat, showing that this species has a broad diet in Atlantic Patagonia, informing about its trophic placement and also suggesting that there are previously unnoticed differences between male and female crab diets. This highlights the importance of monitoring coastal ecosystems, in particular in relation to their invasive species, to be able to detect as early as possible and better understand the negative changes that they can introduce.

## Acknowledgements

We would like to thank Florencia Ríos, Manuela Funes, Juan Pablo Livore, and Maria Martha Mendez for their fieldwork support. We would also like to thank Agustina Ferrando, Cynthia Ibarra, Pablo Macchi and Germán Cheli for expert assessment on taxonomic identifications. We are grateful to two anonymous reviewers for their constructive feedback on the original manuscript version.

## Notes

**Funding** This research was supported by Consejo Nacional de Investigaciones Científicas y Técnicas (CONICET, Argentina) and Sistema Nacional de Datos Genómicos (SNDG, Argentina). Fieldwork campaigns and stable isotope analysis were funded by Fondo para la Investigación Científica y Tecnológica (FonCyT) under the grants PICT 2017-2403, PICT 2018-0903 and PICT 2018-0969. The metabarcoding analysis was funded by the Natural Sciences and Engineering Research Council of Canada and the Canada First Research Excellence Fund.

**Conflicts of interest** The authors declare that they have no conflict of interest.

### Competing Interest Statement

The authors have declared no competing interest.

### Summary of Updates

The new versionis the accepted version in the journal Biological Invasions. The final version includes revised supplementary material.

## REFERENCES

Albaina A, Fox C J, Taylor N, Hunter E, Maillard M, and Taylor MI (2010) A TaqMan real-time PCR based assay targeting plaice (Pleuronectes platessa L.) DNA to detect predation by the brown shrimp (Crangon crangon L.) and the shore crab (Carcinus maenas L.)—assay development and validation. Journal of Experimental Marine Biology and Ecology, 391(1-2), 178–189.

Alekseyenko AV (2016) Multivariate Welch t-test on distances. Bioinformatics, 32(23), 3552–3558.

Arnett RT, and Whelan J (2001) Comparing the diet of cod (Gadus morhua) and grey seals (Halichoerus grypus): an investigation of secondary ingestion. Marine Biological Association of the United Kingdom. Journal of the Marine Biological Association of the United Kingdom, 81(2), 365.

Bertness MD, Crain CM, Silliman BR, Bazterrica MC, Reyna MV, Hildago F, and Farina JK (2006) The community structure of western Atlantic Patagonian rocky shores. Ecological Monographs, 76(3), 439–460.

Bigatti G, and Signorelli J (2018) Marine invertebrate biodiversity from the Argentine Sea, south western Atlantic. ZooKeys, (791), 47.

Boecklen WJ, Yarnes CT, Cook BA, and James AC (2011) On the use of stable isotopes in trophic ecology. Annual Review Of Ecology, Evolution, and Systematics, 42, 411–440.

Boschi EE, Fischbach CE, and Iorio MI (1992) Catálogo ilustrado de los crustáceos estomatópodos y decápodos marinos de Argentina.

Braukmann TW, Ivanova NV, Prosser SW, Elbrecht V, Steinke D, Ratnasingham S, de Waard JR, Sones JE, Zakharov EV, and Hebert PD (2019) Metabarcoding a diverse arthropod mock community. Molecular Ecology Resources, 19(3), 711–727.

Bruno JF, Fridley JD, Bromberg KD, and Bertness MD (2005) Insights into biotic interactions from studies of species invasions. Species invasions: Insights into ecology, evolution, and biogeography, 13–40.

Carlton JT, and Cohen AN (2003) Episodic global dispersal in shallow water marine organisms: the case history of the European shore crabs Carcinus maenas and C. aestuarii. Journal of Biogeography, 30(12), 1809–1820.

Camphuysen CJ, Berrevoets CM, Cremers HJWM, Dekinga A, Dekker R, Ens BJ, van der Have TM, Kats RKH, Kuiken T, Leopold MF, van der Meer J, and Piersma, T (2002) Mass mortality of common eiders (Somateria mollissima) in the Dutch Wadden Sea, winter 1999/2000: starvation in a commercially exploited wetland of international importance. Biological Conservation, 106(3), 303–317.

Canty A, and Ripley B (2021) boot: bootstrap R (S-Plus) Functions. Package version 1.3-27.

Chen RB, Watanabe S, and Yokota M (2004) Feeding habits of an exotic species, the Mediterranean green crab Carcinus aestuarii, in Tokyo Bay. Fisheries Science, 70(3), 430–435.

Clare EL (2014) Molecular detection of trophic interactions: emerging trends, distinct advantages, significant considerations and conservation applications. Evolutionary Applications, 7(9), 1144–1157.

Clavero M, and García-Berthou E (2005) Invasive species are a leading cause of animal extinctions. Trends in Ecology and Evolution, 20(3), 110.

Cohen AN, Carlton JT, and Fountain MC. (1995) Introduction, dispersal and potential impacts of the green crab Carcinus maenas in San Francisco Bay, California. Marine Biology, 122(2), 225–237.

Compson ZG, Monk WA, Hayden B, Bush A, O’Malley Z, Hajibabaei M, Porter TM, Wright MTG, Baker CJO, Sadnan Al Manir M, Curry RA, and Baird DJ (2019) Network-based biomonitoring: exploring freshwater food webs with stable isotope analysis and DNA metabarcoding. Frontiers in Ecology and Evolution, 7: 395. doi: 10.3389/fevo.

Dare PJ, and Edwards DB (1981) Underwater television observations on the intertidal movements of shore crabs, Carcinus maenas, across a mudflat. Journal of the Marine biological Association of the United Kingdom, 61(1), 107–116.

Darling JA (2011) Interspecific hybridization and mitochondrial introgression in invasive Carcinus shore crabs. Plos One, 6(3), e17828.

David P, Thebault E, Anneville O, Duyck PF, Chapuis E, and Loeuille N (2017) Impacts of invasive species on food webs: a review of empirical data. In Advances in Ecological Research (Vol. 56, pp. 1–60). Academic Press.

De Lecea AM, Smit AJ, and Fennessy ST (2011) The effects of freeze/thaw periods and drying methods on isotopic and elemental carbon and nitrogen in marine organisms, raising questions on sample preparation. Rapid Communications in Mass Spectrometry, 25(23), 3640–3649

Deagle BE, Thomas AC, McInnes JC, Clarke LJ, Vesterinen EJ, Clare EL, Kartzinel TR, and Eveson JP (2019) Counting with DNA in metabarcoding studies: How should we convert sequence reads to dietary data? Molecular Ecology, 28(2), 391–406.

DeGraaf JD, and Tyrrell MC (2004) Comparison of the feeding rates of two introduced crab species, Carcinus maenas and Hemigrapsus sanguineus, on the blue mussel, Mytilus edulis. Northeastern Naturalist, 11(2), 163–167.

Doherty TS, Glen AS, Nimmo DG, Ritchie EG, and Dickman CR (2016) Invasive predators and global biodiversity loss. Proceedings of the National Academy of Sciences, 113(40), 11261–11265.

Ebling FJ, Kitching JA, Muntz L, and Taylor CM (1964) Experimental observations of the destruction of Mytilus edulis and Nucella lapillus by crabs. Journal of Animal Ecology, 33, 73–82.

Efron B, and Tibshirani RJ (1993) An introduction to the bootstrap Chapman & Hall. New York, 436.

Elbrecht V, Braukmann TW, Ivanova NV, Prosser SW, Hajibabaei M, Wright M, Zakharov EV, Hebert PDN, and Steinke D (2019) Validation of COI metabarcoding primers for terrestrial arthropods. PeerJ, 7, e7745.

Elbrecht V, and Steinke D (2019) Scaling up DNA metabarcoding for freshwater macrozoobenthos monitoring. Freshwater Biology, 64(2), 380–387.

Elbrecht V, Vamos EE, Meissner K, Aroviita J, and Leese F (2017) Assessing strengths and weaknesses of DNA metabarcoding-based macroinvertebrate identification for routine stream monitoring. Methods in Ecology and Evolution, 8(10), 1265–1275.

Elbrecht V, and Leese F (2017) Validation and development of COI metabarcoding primers for freshwater macroinvertebrate bioassessment. Frontiers in Environmental Science, 5, 11.

Elner R.W (1981) Diet of green crab Carcinus maenas (L.) from Port Hebert, southwestern Nova Scotia. J. Shellfish Res. 1, 89–94.

Elner RW (1980) The influence of temperature, sex and chela size in the foraging strategy of the shore crab, Carcinus maenas (L.). Marine and Freshwater Behaviour and Physiology, 7(1), 15–24.

Fauchald K (1977) The polychaete worms. Definitions and keys to the orders, families and genera. Natural History Museum of Los Angeles County, Science Series.

Fridley JD, Stachowicz JJ, Naeem S, Sax DF, Seabloom EW, Smith MD, Stohlgren TJ, Tilman D, and Holle BV (2007) The invasion paradox: reconciling pattern and process in species invasions. Ecology, 88(1), 3–17.

Galloway AW, Brett MT, Holtgrieve GW, Ward EJ, Ballantyne AP, Burns CW, Kainz MJ, Müller-Navarra DC, Persson J, Ravet JL, Strandberg U, Taipale SJ, and Alhgren G (2015) A fatty acid based Bayesian approach for inferring diet in aquatic consumers. PloS one, 10(6), e0129723.

García-Robledo C, Erickson DL, Staines CL, Erwin TL, and Kress WJ (2013) Tropical plant–herbivore networks: reconstructing species interactions using DNA barcodes. PLoS One, 8(1), e52967.

Gehrels H, Tummon Flynn P, Cox R, and Quijón PA (2017) Effects of habitat complexity on cannibalism rates in European green crabs (Carcinus maenas Linnaeus, 1758). Marine Ecology, 38(5), e12448

Gosztonyi AE, and Kuba L (1996) Atlas de huesos craneales y de la cintura escapular de peces costeros patagónicos. Informe Técnico: Plan de Manejo integrado de la zona costera patagónica. Fundación Patagonia Natural.

Grosholz ED, and Ruiz GM (1995) Spread and potential impact of the recently introduced European green crab, Carcinus maenas, in central California. Marine Biology, 122(2), 239–247.

Hamidi B, Wallace K, Vasu C, and Alekseyenko AV (2019) W d*$ W_ {d}^{*} $-test: robust distance-based multivariate analysis of variance. Microbiome, 7(1), 1–9.

Hayden B, Harrod C, and Kahilainen KK (2014) Dual fuels: Intra-annual variation in the relative importance of benthic and pelagic resources to maintenance, growth and reproduction in a generalist salmonid fish. Journal of Animal Ecology, 83(6), 1501–1512.

Hebert PD, Cywinska A, Ball SL, and Dewaard JR (2003) Biological identifications through DNA barcodes. Proceedings of the Royal Society of London. Series B: Biological Sciences, 270(1512), 313–321.

Hidalgo FJ, Silliman BR, Bazterrica MC, and Bertness MD (2007) Predation on the rocky shores of Patagonia, Argentina. Estuaries and Coasts, 30(5), 886–894.

Howard BR, Barrios-O’Neill D, Alexander ME, Dick JT, Therriault TW, Robinson TB, and Côté IM (2018) Functional responses of a cosmopolitan invader demonstrate intraspecific variability in consumer-resource dynamics. PeerJ, 6, e5634.

Hui C, Richardson DM, Landi P, Minoarivelo HO, Garnas J, and Roy HE (2016) Defining invasiveness and invasibility in ecological networks. Biological Invasions, 18(4), 971–983.

Hussey NE, MacNeil MA, McMeans BC, Olin JA, Dudley SF, Cliff G, Wintner SP, Fennessy ST, and Fisk AT (2014) Rescaling the trophic structure of marine food webs. Ecology letters, 17(2), 239–250.

Hyndes G, Hanson C, Vanderklift M (2013) The magnitude of spatial and temporal variation in δ^15^N and δ^13^C differs between taxonomic groups: implications for food web studies. Estuar. Coast. Shelf Sci. 119, 176–187.

Hyslop EJ (1980) Stomach contents analysis—a review of methods and their application. Journal of Fish Biology, 17(4), 411–429.

Jackson AL, Inger R, Parnell AC, and Bearhop S (2011) Comparing isotopic niche widths among and within communities: SIBER–Stable Isotope Bayesian Ellipses in R. Journal of Animal Ecology, 80(3), 595–602.

Klassen GJ, and Locke A (2007) A biological synopsis of the European green crab, Carcinus maenas. Moncton, New Brunswick, Canada: Fisheries and Oceans Canada.

Koen Alonso MK, Crespo EA, García NA, Pedraza SN, Mariotti PA, and Mora NJ (2002) Fishery and ontogenetic driven changes in the diet of the spiny dogfish, Squalus acanthias, in Patagonian waters, Argentina. Environmental Biology of Fishes, 63(2), 193–202.

Lefebvre S, and Dubois S (2016) The stony road to understand isotopic enrichment and turnover rates: insight into the metabolic part. Vie Et Milieu-life And Environment, 66(3-4), 305–314.

Leignel VSJH, Stillman JH, Baringou S, Thabet R, and Metais I (2014) Overview on theEuropean green crab Carcinus spp. (Portunidae, Decapoda), one of the most famous marine invaders and ecotoxicological models. Environmental Science and Pollution Research, 21(15), 9129–9144.

Mackey AP, Hyndes GA, Carvalho MC, and Eyre BD (2015) Physical and biogeochemical correlates of spatio-temporal variation in the δ^13^C of marine macroalgae. Estuarine, Coastal and Shelf Science, 157, 7–18.

McDonald PS, Jensen GC, and Armstrong DA (2001) The competitive and predatory impacts of the nonindigenous crab Carcinus maenas (L.) on early benthic phase Dungeness crab Cancer magister Dana. Journal of Experimental Marine Biology and Ecology, 258(1), 39–54.

Majdi, N, Hette-Tronquart N, Auclair E, Bec A, Chouvelon T, Cognie B, Danger M, Decottignies P, Dessier A, Desvilettes C, Dubois S, Dupuy C, Fritsch C, Gaucherel C, Hedde M, Jabot F, Lefebvre S, Marzloff M, Pey B, Peyrard N, Powolny T, Sabbadin R, Thebault E, & Perga ME. (2018) There’s no harm in having too much: A comprehensive toolbox of methods in trophic ecology. Food webs, 17, e00100.

McMurdie PJ, and Holmes S (2013) Phyloseq: An R package for reproducible interactive analysis and graphics of microbiome census data. PLoS ONE 8(4):e61217.

Mendez MM, Livore JP, Márquez F, Gotlieb E, and Bigatti G (2019) Drástica pérdida de cobertura en mejillinares del golfo nuevo.3° Congreso Argentino de Malacología. 4 al 6 de diciembre de 2019. Bahía Blanca, Argentina.

Miron G, Audet D, Landry T, and Moriyasu M (2005) Predation potential of the invasive green crab (Carcinus maenas) and other common predators on commercial bivalve species found on Prince Edward Island. Journal of Shellfish Research, 24(2), 579–586.

Mooney CZ, and Duvall R (1993) Bootstrapping: A nonparametric approach to statistical inference (No. 95).

Nakahara F, Ando H, Ito H, Murakami A, Morimoto N, Yamasaki M, Takayanagi A, and Isagi Y (2015) The applicability of DNA barcoding for dietary analysis of sika deer. DNA Barcodes, 3(1).

Nielsen JM, Clare EL, Hayden B, Brett MT, and Kratina P (2018) Diet tracing in ecology: Method comparison and selection. Methods in Ecology and Evolution, 9(2), 278–291.

Oksanen J, Blanchet, FG, Kindt R, Legendre P, O’hara RB, Simpson GL, Solymos P, Henry M, Szoecs S, Szoecs E, and Wagner H (2019) Vegan: community ecology package. R package version 2.5-6. URL http://CRAN.R-project.org/package=vegan.

Pardal M, Baeta A, Marques J, and Cabral H (2006) Feeding ecology of the green crab, Carcinus maenas (L., 1758) in a temperate estuary, Portugal. Crustaceana, 79(10), 1181–1193.

Parker PL, Anderson RK, and Lawrence A (1989) A δ^13^C and δ^15^N tracer study of nutrition in aquaculture: Penaeus vannamei in a pond growout system. In Stable isotopes in ecological research (pp. 288–303). Springer, New York, NY.

Philippsen JS, and Benedito E (2013) Discrimination factor in the trophic ecology of fishes: a review about sources of variation and methods to obtain it. Oecologia Australis, 17(2), 205–2016.

Pielou EC (1966) The measurement of diversity in different types of biological collections. Journal of Theoretical Biology, 13, 131–144.

Pinnegar JK, and Polunin NVC (1999) Differential fractionation of δ^13^C and δ^15^N among fish tissues: implications for the study of trophic interactions. Functional Ecology, 13(2), 225–231.

Pompanon F, Deagle BE, Symondson WO, Brown DS, Jarman SN, and Taberlet P (2012) Who is eating what: diet assessment using next generation sequencing. Molecular Ecology, 21(8), 1931–1950.

Post, D M (2002) Using stable isotopes to estimate trophic position: models, methods, and assumptions. Ecology, 83(3), 703–718.

Quezada-Romegialli C, Jackson AL, Hayden B, Kahilainen KK, Lopes C, and Harrod C (2018) tRophicPosition, an R package for the Bayesian estimation of trophic position from consumer stable isotope ratios. Methods in Ecology and Evolution, 9(6), 1592–1599.

R Core Team (2020) R: A language and environment for statistical computing. R Foundation for Statistical Computing, Vienna, Austria. URL https://www.R-project.org/.

Ratnasingham S (2019) mBRAVE: The Multiplex Barcode Research And Visualization Environment. Biodiversity Information Science and Standards, 3, e37986.

Ratnasingham S, and Hebert PD (2007) BOLD: The Barcode of Life Data System (http://www.barcodinglife.org). Molecular Ecology Notes, 7(3), 355–364.

Rechimont ME, Galvan DE, Sueiro MC, Casas G, Piriz ML, Diez ME, Primost M, Zabala MS, Márquez F, Brogger M, Alfaya JF, and Bigatti G (2013) Benthic diversity and assemblage structure of a north Patagonian rocky shore: a monitoring legacy of the NaGISA project. Marine Biological Association of the United Kingdom. Journal of the Marine Biological Association of the United Kingdom, 93(8), 2049.

Reid, DG, Abelló P, Kaiser, MJ, and Warman CG (1997) Carapace colour, inter-moult duration and the behavioural and physiological ecology of the shore crab Carcinus maenas. Estuarine, Coastal and Shelf Science, 44(2), 203–211.

Rodriguez LF (2006) Can invasive species facilitate native species? Evidence of how, when, and why these impacts occur. Biol Invasions 8: 927−939

Romanuk TN, Zhou Y, Valdovinos FS, and Martinez ND (2017) Robustness trade-offs in model food webs: invasion probability decreases while invasion consequences increase with connectance. In Advances in Ecological Research (Vol. 56, pp. 263–291). Academic Press.

Rossong MA, Quijón PA, Snelgrove PV, Barrett TJ, McKenzie CH, and Locke A (2012) Regional differences in foraging behaviour of invasive green crab (Carcinus maenas) populations in Atlantic Canada. Biological Invasions, 14(3), 659–669.

Rudnick D, and Resh V (2005) Stable isotopes, mesocosms and gut content analysis demonstrate trophic differences in two invasive decapod crustacea. Freshwater Biology, 50(8), 1323–1336

Russel J (2020) MicEco: Various functions for microbial community data. R package version 0.9.9.

Schwindt E, Carlton JT, Orensanz JM, Scarabino F, and Bortolus A (2020) Past and future of the marine bioinvasions along the Southwestern Atlantic. Aquatic Invasions, 15(1).

Sigurdsson GM, and Rochette R (2013) Predation by green crab and sand shrimp on settling and recently settled American lobster postlarvae. Journal of Crustacean Biology, 33, 10–14

Silliman BR, Bertness MD, Altieri AH, Griffin JN, Bazterrica MC, Hidalgo FJ, Crain CM, and Reyna MV (2011) Whole-community facilitation regulates biodiversity on Patagonian rocky shores. PLoS One, 6(10), e24502.

Singh R (1991) Natural Diet of, and Shelter-Use by, the Green Crab, Carcinus maenas (L.) in Nova Scotia. Master’s Thesis, University of New Brunswick, Fredericton, NB, Canada.

Smith PJ, McVeagh SM, Allain V, and Sanchez C (2005) DNA identification of gut contents of large pelagic fishes. Journal of Fish Biology, 67(4), 1178–1183.

Spivak ED, Farías NE, Ocampo EH, Lovrich GA, and Luppi TA (2019) Annotated catalogue and bibliography of marine and estuarine shrimps, lobsters, crabs and their allies (Crustacea: Decapoda) of Argentina and Uruguay (Southwestern Atlantic Ocean). Frente Marítimo, 26, 1–164.

Spooner EH, Coleman RA, and Attrill MJ (2007) Sex differences in body morphology and multitrophic interactions involving the foraging behaviour of the crab Carcinus maenas. Marine Ecology, 28(3), 394–403.

Stock BC, Jackson AL, Ward EJ, Parnell AC, Phillips DL, and Semmens BX (2018) Analyzing mixing systems using a new generation of Bayesian tracer mixing models. PeerJ, 6, e5096.

Taberlet P, Bonin A, Coissac E, and Zinger L (2018) Environmental DNA: For biodiversity research and monitoring. Oxford University Press.

Taberlet P, Coissac E, Pompanon F, Brochmann C, and Willerslev E (2012) Towards next-generation biodiversity assessment using DNA metabarcoding. Molecular ecology, 21(8), 2045–2050.

Thomsen MS, Byers JE, Schiel DR, Bruno JF, Olden JD, Wernberg T, and Silliman BR (2014) Impacts of marine invaders on biodiversity depend on trophic position and functional similarity.Marine Ecology Progress Series, 495, 39–47.

Thomsen MS, Olden JD, Wernberg T, Griffin JN, and Silliman BR (2011a) A broad framework to organize and compare ecological invasion impacts. Environmental Research, 111(7), 899–908.

Thomsen MS, Wernberg T, Olden JD, Griffin JN, and Silliman BR (2011b) A framework to study the context-dependent impacts of marine invasions. Journal of Experimental Marine Biology and Ecology, 400(1-2), 322–327.

Thomsen MS, Wernberg T, Altieri A, Tuya F, Gulbransen D, McGlathery KJ, Holmer M, and Silliman BR (2010) Habitat cascades: the conceptual context and global relevance of facilitation cascades via habitat formation and modification. Integrative and Comparative Biology, 50(2), 158–175.

Tilman D, Isbell F, and Cowles JM (2014) Biodiversity and ecosystem functioning. Annual Review of Ecology, Evolution, and Systematics, 45, 471–493.

Tirasin EM, and Jørgensen T (1999) An evaluation of the precision of diet description. Marine Ecology Progress Series, 182, 243–252.

Torchin ME, Lafferty KD, and Kuris AM (1996) Infestation of an introduced host, the European green crab, Carcinus maenas, by a symbiotic nemertean egg predator, Carcinonemertes epialti. The Journal of Parasitology, 449–453.

Torres PJ, and González-Pisani X (2016) Primer registro del cangrejo verde, Carcinus maenas (Linnaeus, 1758), en Golfo Nuevo, Argentina: un nuevo límite norte de distribución en costas patagónicas. Ecología Austral, 26(2), 134–137.

Thalinger B, Oehm J, Zeisler C, Vorhauser J, and Traugott M (2018) Sex-specific prey partitioning in breeding piscivorous birds examined via a novel, noninvasive approach. Ecology and Evolution, 8(17), 8985–8998.

Traugott M, Thalinger B, Wallinger C, and Sint D (2021) Fish as predators and prey: DNA-based assessment of their role in food webs. Journal of Fish Biology. https://doi.org/10.1111/jfb.14400

Traugott M, Kamenova S, Ruess L, Seeber J, and Plantegenest M (2013) Empirically characterising trophic networks: what emerging DNA-based methods, stable isotope and fatty acid analyses can offer. In Advances in Ecological Research (Vol. 49, pp. 177–224). Academic Press.

Unger SD, Williams LA, Diaz L, and Jachowski CB (2020) DNA barcoding to assess diet of larval eastern hellbenders in North Carolina. Food Webs, 22, e00134.

Whitaker MRL, Baker CCM, Salzman SM, Martins DJ, Pierce NE (2019) Combining stable isotope analysis with DNA metabarcoding improves inferences of trophic ecology. PLoS ONE 14(7): e0219070. https://doi.org/10.1371/journal.pone.0219070

White EM, Wilson JC, and Clarke AR (2006) Biotic indirect effects: a neglected concept in invasion biology. Diversity and Distributions, 12(4), 443–455.

Wilcox M, and Rochette R (2015) Does claw morphology of the green crab Carcinus maenas vary in relation to its diet on rocky versus fine-sediment shores of southwest New Brunswick, Bay of Fundy, Canada? Journal of Experimental Marine Biology and Ecology, 465, 121–129.

Yokoyama H, Tamaki A, Harada K, Shimoda K, Koyama K, and Ishihi Y (2005) Variability of diet-tissue isotopic fractionation in estuarine macrobenthos. Marine Ecology Progress Series, 296, 115–128.

Yorio P, Suárez N, Kasinsky T, Pollicelli M, Ibarra C, and Gatto, A (2020) The introduced green crab (Carcinus maenas) as a novel food resource for the opportunistic kelp gull (Larus dominicanus) in Argentine Patagonia. Aquatic Invasions, 15(1), 140–159.

Young AM, and Elliott JA (2020) Life History and Population Dynamics of Green Crabs (Carcinus maenas). Fishes, 5(1), 4.

Zacharia P, Abdurahiman K, and Mohamed K (2004) Methods of stomach content analysis of fishes. Winter School on Towards Ecosystem Based Management of Marine Fisheries— Building Mass Balance Trophic and Simulation Models, 1, 148–158.

