## Supplemental material 2 for "Metabarcoding, direct stomach observation and stable isotope analysis reveal a highly diverse diet for the invasive green crab in Atlantic Patagonia"

Electronic Supplemental Material
Online Resource 2

**Title:**

### Metabarcoding, direct stomach observation and stable isotope analysis reveal a highly diverse diet for the invasive green crab in Atlantic Patagonia


**Authors--**

Georgina Cordone[1], Mariana Lozada[2], Elisabet Vilacoba[3], Bettina Thalinger[4], Gregorio Bigatti[5,6], Darío A. Lijtmaer[3], Dirk Steinke[4], David E. Galván[1]*

[1] Centro para el Estudio de Sistemas Marinos (CESIMAR), Consejo Nacional de Investigaciones Científicas y Técnicas (CONICET), Edificio CCT CONICET-CENPAT, Bvd. Brown 2915, (U9120) Puerto Madryn, Chubut, Argentina.

[2] Laboratorio de Microbiología Ambiental (IBIOMAR), Consejo Nacional de Investigaciones Científicas y Técnicas (CONICET), Edificio CCT CONICET-CENPAT, Bvd. Brown 2915, (U9120) Puerto Madryn, Chubut, Argentina.

[3] División Ornitología, Museo Argentino de Ciencias Naturales “Bernardino Rivadavia” MACN-CONICET. Av. Ángel Gallardo 470 (C1405) Ciudad Autónoma de Buenos Aires, Buenos Aires, Argentina.

[4] Centre for Biodiversity Genomics,University of Guelph, 50 Stone Rd E (N1G2W1), Guelph, Ontario, Canada.

[5] Laboratorio de Reproducción y Biología Integrativa de Invertebrados Marinos, (LARBIM, IBIOMAR). Consejo Nacional de Investigaciones Científicas y Técnicas (CONICET), Edificio CCT CONICET-CENPAT, Bvd. Brown 2915, (U9120) Puerto Madryn, Chubut, Argentina.

[6] Universidad Espíritu Santo, Ecuador

#### 2. Materials and Methods

##### 2.3 Metabarcoding analysis of gut content

*Sequence pre-processing*

Sequence pre-processing was carried out using JAMP (https://github.com/VascoElbrecht/JAMP ). Sequences were demultiplexed and forward and reverse reads paired-end merged. Primer sequences were removed, before setting a minimum and a maximum sequence length threshold that allowed further processing. As we expected a highly diverse set of taxa, sequences between 380bp and 440 bp remained in the dataset (+/- 30bp of expected fragment length). The resulting files were processed in two ways:

i. **The reads were quality filtered and taxonomy assigned using mBRAVE (BOLD)**
ii. **Quality filtering was carried out in JAMP and the resulting haplotypes were blasted against the NCBI nucleotide database**

*Quality filtering and assignment using mBRAVE and BOLD*

Multiplex Barcode Research And Visualization Environment of the Barcode of Life platform (mBRAVE/BOLD, http://www.mbrave.net/) was used in this case for comparison of sequences against a closed-reference, curated database. In this case, reads were merged, trimmed to 425 bp, reads shorter than 100 bp were eliminated, and reads with more than 10% of bases with low (<20) quality values (QV), or with more than 1% of Ultra Low (<10) QV were eliminated.

The reads are not clustered because mBRAVE uses a closed reference approach for assignment to BINs in the reference database. Sequences are matched to selected BIN (Barcode

Index Numbers) databases retrieved from BOLD using the *refined single linkage algorithm* (*RESL*) [1]. We used 3% as the clustering threshold, and the minimum OTU Size considered was 5.

The system mBRAVE accesses all BOLD data (including private datasets) to retrieve taxonomic information. The following databases were used to contrast the data:

- SYS-CRLBACTERIA: Bacteria COI , 9746 sequences, 2113 BINs, 2066 species, 12-Jul-2020
- SYS-CRLNONARTHINVERT: Non-Arthropoda Invertebrates, 77197 sequences, 46519 BINs, 31663 species, 12-Jul-2020
- SYS-CRLNONINSECTARTH: Non-Insect Arthropoda, 84884 sequences, 63733 BINs, 25255 species, 12-Jul-2020
- SYS-CRLINSECTA: Insecta, 708485 sequences, 516810 BINs, 211284 species, 12-Jul-2020

*Quality filtering and denoising in JAMP, blasted against the NCBI nucleotide database*

In JAMP, common haplotypes are prefiltered, extracted using a denoising module (command denoise(), unoise3 algorithm, https://www.drive5.com/usearch/manual/cmd_unoise3.html) to perform error-correction as well as chimera detection and removal in amplicon reads, followed by OTU clustering at 97% identity plus abundance-based filtering.

In this case, paired end reads were merged, and quality filtering was applied using the command *U_max_ee* with an expected error value set to 1. All sequences with less than 5 reads **or** contributing less than 0.1% to the total library **or** OTUs with a relative abundance below 1% were removed during the Denoising process. (For details on denoising see https://github.com/VascoElbrecht/JAMP/wiki/5.-Denoising-quick-guide). The obtained haplotypes were blasted against the NCBI nucleotide database (nt) as of 7th Feb. 2020.

**References**

**[1]** Ratnasingham S, and Hebert PDN (2013) A DNA-Based Registry for All Animal Species: The Barcode Index Number (BIN) System. PLOS ONE 8:e66213. https://doi.org/10.1371/journal.pone.0066213

#### 2.5 Stable isotope analysis

*Equations for TL estimations*

Eq. A1, Eq. A2 and Eq. A3 correspond to the dual baseline Bayesian mixing model proposed by Quezada-Romegialli et al., (2018), where $\delta^{15}N_{b1}$ , $\delta^{13}C_{b1}$ , $\delta^{15}N_{b2}$ and $\delta^{13}C_{b2}$ refer to the $\delta^{15}N$ and $\delta^{13}C$ values of baselines 1 and 2 and α is the proportion of N derived from baseline 1 but including the trophic discrimination factor for carbon (ΔC) (Post 2002; Quezada-Romegialli et al., 2018).

$$\delta^{15}N_c = \Delta(TP+\lambda) + \alpha(\delta^{15}N_{b1} + \delta^{15}N_{b2}) - \delta^{15}N_{b2} \quad \text{(eq. 1)}$$

$$\delta^{13}C_c = \delta^{13}C_{b1}\alpha + \delta^{13}C_{b2}(1-\alpha) \quad \text{(eq. 2)}$$

$$\alpha = {}^{(\delta^{13}C_{b2} - (\delta^{13}C_c + \Delta C))/(TP-\lambda)}\Big/_{\delta^{13}C_{b2} + \delta^{13}C_{b1}} \quad \text{(eq. 3)}$$

### 3. Results

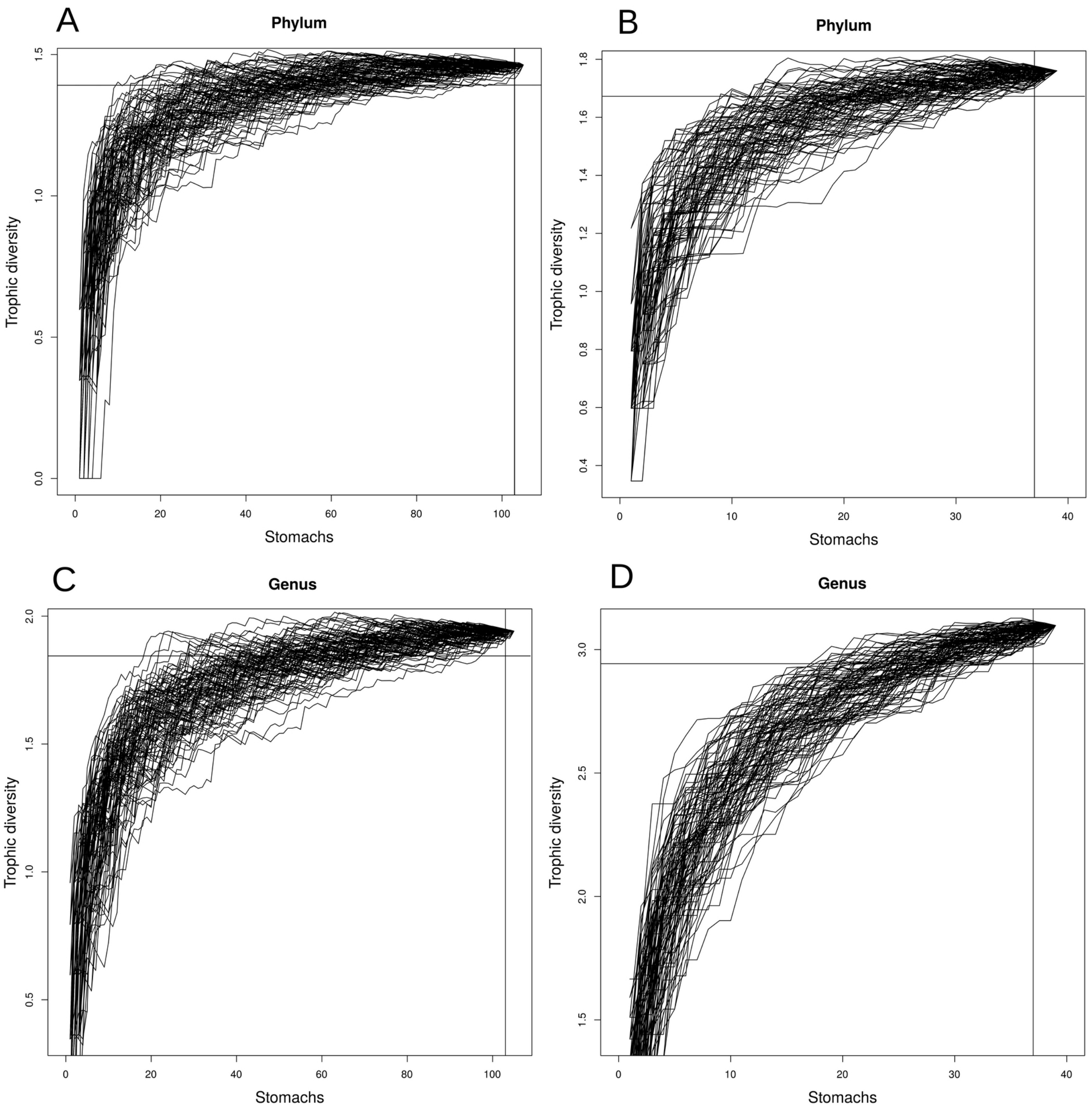


**Figure S1.** Trophic diversity accumulation curves for prey items at two taxonomic classification levels for data obtained by **A and C)** visual (phylum and genus respectively) and **B and D)** metabarcoding analyses (phylum and genus). Horizontal lines show Brillouin diversity index (Hz) values (Hz ± 0.05 Hz) and the vertical line shows (n-2) values where n is the number of samples with at least one prey item (i.e. no empty stomachs).

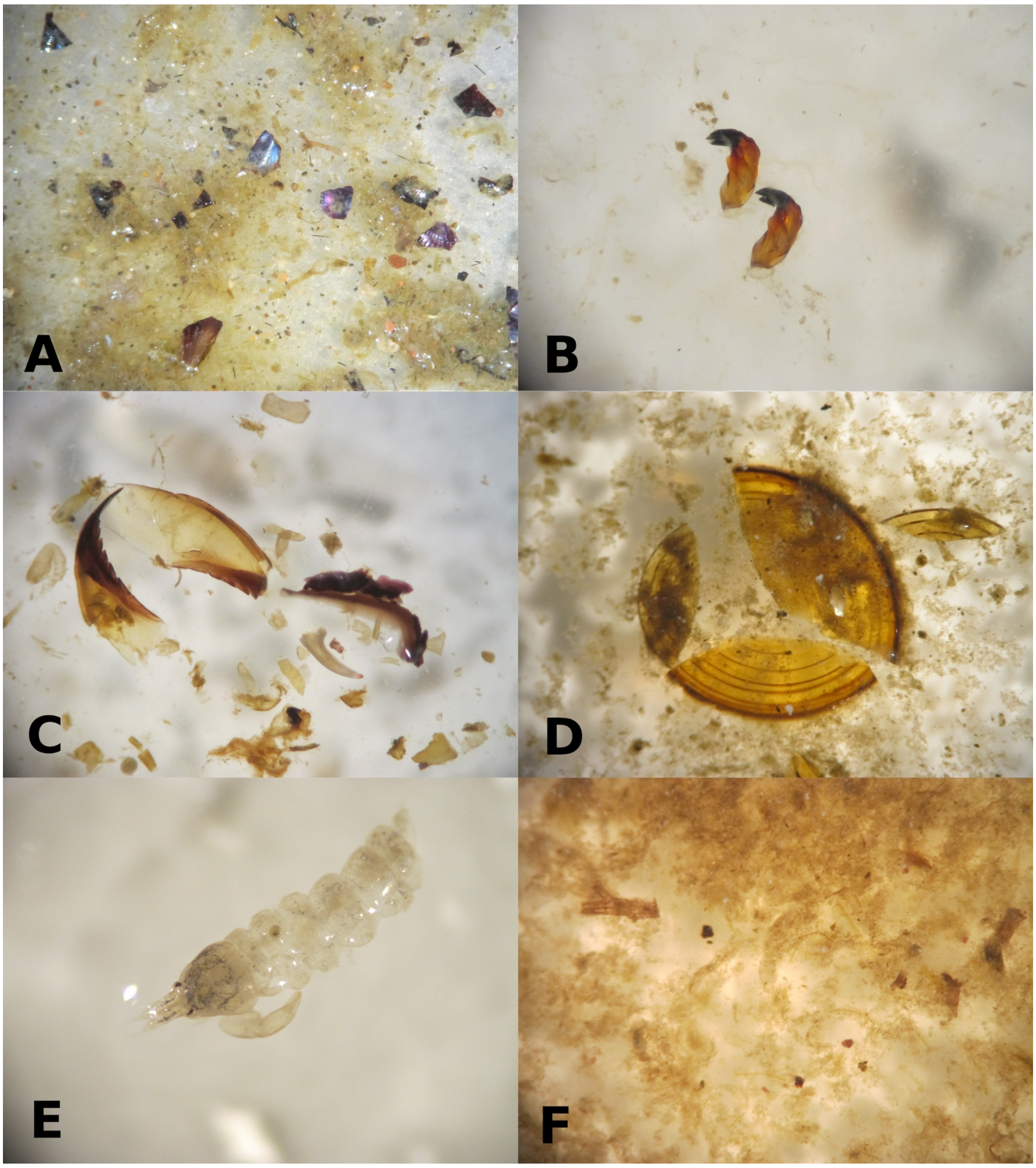


**Figure S2.** Prey items found in stomach contents of green crabs: **A)** Mytilids, **B)** Chiton radula, **C)** Polychaeta (Nereidae) jaws, **D)** Gastropod opercule (*Tegula patagonica*), **E)** Tanaidacea remains (*Tanais dulongii*) and **F)** unidentified material. Photo credits: Georgina Cordone.

**Table S1.** Taxonomic assignment of COI sequences based on the BOLD database as implemented in mBRAVE system (mBRAVE/BOLD dataset). The lowest taxonomic rank reached based on local records and/or BOLD's BIN information is marked in bold. Items not considered prey are depicted in grey. BIN: name of the reference Barcode Index Numbers (BINs) as defined by BOLD Systems (www.boldsystems.org). av_dist_BIN(%): average percent genetic distance among members of the reference BIN. max_dist_BIN(%): maximum percent distance among members of the reference BIN. species_bin: species recorded in the BIN. location_bin: location where the members of the BIN were recorded. nsequences: number of sequences. nsamples: number of samples where the item was observed. **See in Online Resource 3.**

**Table S2.** Taxonomic assignment of COI sequences based on blast results against NCBI nucleotide (nt) database. The first tab "otus_ncbi_new_taxa" shows only members of families not appearing in BOLD, while the second tab "all_otus_ncbi_blast_firstmatch" shows all OTUs with their respective first blast match . The lowest taxonomic rank based on percent identity to the first match is shown in bold. qseqid: query sequence identifier; sseqid: subject (blast hit) sequence identifier; pident: pairwise identity(%); length: alignment length (bases); qcovs: alignment coverage based on query mismatch: mismatches in alignment; gapopen: gap openings as defined by blast; qstart: query start position; qend: query end position; sstart: subject start position; send: subject end position; evalue: E-value bitscore: score of the alignment as defined by blast; staxids: subject taxonomic identifiers in NCBI Taxonomy; OTU: OTU number. **See in Online Resource 4.**

**Table S3.** Pairwise comparisons of trophic position (*TP*) and alpha (*α*) posterior estimates for female and males green crab at Patagonia Argentina. This matrix was constructed using the 'pairwiseComparisons' function of the R package 'tRophic Position' (version 0.7.7). There are not ** as none of the probabilities is higher than 0.95 (i.e. significant).

| | TP females | TP males | *α* females | *α* males |
|---|---|---|---|---|
| TP females | 0.000 | 0.278 | - | - |
| TP males | 0.722 | 0.000 | - | - |
| *α* females | - | - | 0.000 | 0.273 |
| *α* males | - | - | 0.727 | 0.000 |

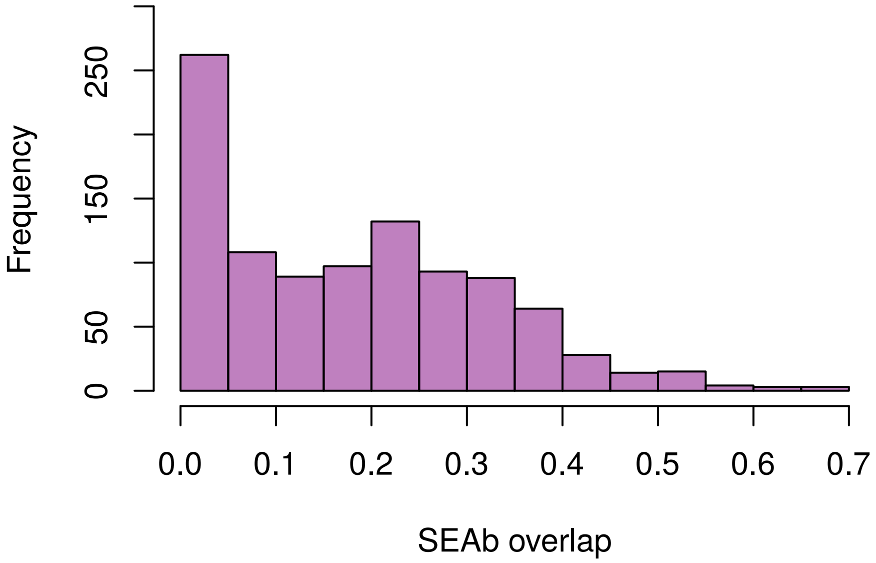


**Figure S3.** Histogram showing the overlap between a sub-sample of 1000 simulated standard ellipse area by Bayesian methods (SEAb) for females and males green crab.

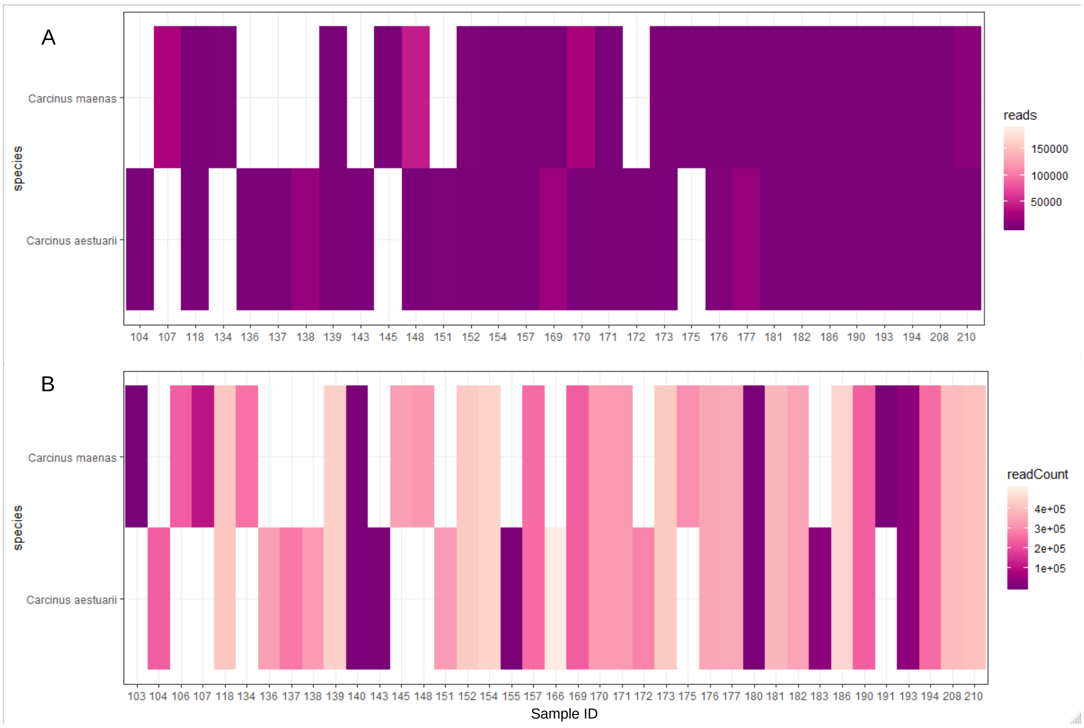


**Figure S4.** Distribution of consumer sequences amongst samples. Both approaches detected sequences of *C. maenas* and *C. aestuarii* in most samples, see their read counts per sample (x-axis) **A**: in the NCBI data and **B**: BOLD data.
